# Conserved residues in the wheat (*Triticum aestivum*) NAM-A1 NAC domain are required for protein binding and when mutated lead to delayed peduncle and flag leaf senescence

**DOI:** 10.1101/573881

**Authors:** Sophie A. Harrington, Lauren E. Overend, Nicolas Cobo, Philippa Borrill, Cristobal Uauy

## Abstract

**Background:** NAC transcription factors contain five highly conserved subdomains which are required for protein dimerisation and DNA binding. Few residues within these subdomains have been identified as essential for protein function, and fewer still have been shown to be of biological relevance *in planta*. Here we use a positive regulator of senescence in wheat, *NAM-A1*, to test the impact of missense mutations at specific, highly conserved residues of the NAC domain on protein function.

**Results:** We identified missense mutations in five highly conserved residues of the NAC domain of *NAM-A1* in a tetraploid TILLING population. TILLING lines containing these mutations, alongside synonymous and non-conserved mutation controls, were grown under glasshouse conditions and scored for senescence. Four of the five mutations showed a significant and consistent delay in peduncle senescence but had no consistent effects on flag leaf senescence. All four mutant alleles with the delayed senescence phenotype also lost the ability to interact with the homoeolog NAM-B1 in a yeast two-hybrid assay. Two of these residues were previously shown to be involved in NAC domain function in Arabidopsis, suggesting conservation of residue function between species. Three of these four alleles led to an attenuated cell death response compared to wild-type *NAM-A1* when transiently over-expressed in *Nicotiana benthamiana*. One of these mutations was further tested under field conditions, in which there was a significant and consistent delay in both peduncle and leaf senescence.

**Conclusions:** We combined field and glasshouse studies of a series of mutant alleles with biochemical analyses to identify four residues of the NAC domain which are required for *NAM-A1* function and protein interaction. We show that mutations in these residues lead to a gradient of phenotypes, raising the possibility of developing allelic series of mutations for traits of agronomic importance. We also show that mutations in *NAM-A1* more severely impact peduncle senescence, compared to the more commonly studied flag leaf senescence, highlighting this as an area deserving of further study. The results from this integrated approach provide strong evidence that conserved residues within the functional domains of NAC transcription factors have biological significance *in planta*.

## Background

NAC transcription factors are a large family of plant-specific transcription factors (TFs). NACs regulate a broad set of biological processes, including many fundamental to development such as lateral root formation, shoot apical meristem development, and leaf senescence (1–5). They also regulate a wide range of abiotic and biotic stress responses (6–8). While much of the earliest research was carried out in model species such as *Arabidopsis thaliana*, increasingly, research in crop species has found NAC TFs to be essential regulators of a variety of agronomically-relevant traits. In wheat, for example, NAC TFs are involved in regulating abiotic and biotic stress responses (9–18), as well as traits involved in determining grain quality (5, 19).

Characteristic of the NAC transcription factors is a highly conserved NAC domain at the N-terminus of the protein, followed by a non-conserved, intrinsically disordered region at the C-terminus (3, 20). The NAC domain consists of five subdomains (subdomains i to v) that are themselves highly conserved across the plant kingdom, and which have critical functional roles (20–22). Early studies in Arabidopsis demonstrated that NAC TFs bind DNA as homo- or hetero-dimers, and that this dimerization is required to stabilise the DNA binding (23, 24). Crystallization of the NAC domain in a homo-dimer form localised the protein dimerisation interface to subdomains i and ii (20). Later, crystal structures of the NAC dimer binding to DNA identified residues in subdomains iii, iv, and v located at the DNA-protein interaction interface (21).

Despite their importance, it is still unclear to what extent residues within the NAC domain share common functions across different NAC transcription factors, and indeed across different species. Early work showed that mutation of a specific residue in subdomain iii of the Arabidopsis ANAC019 NAC domain leads to a loss of DNA binding *in vitro* (Figure 1A) (24). Further investigation identified a pair of residues essential for protein dimerization in ANAC019 (Figure 1A, subdomain i) (24). More recently, various residues within the ANAC019 NAC domain have been predicted to have a role in pH-dependent stabilisation of the NAC domain (subdomain i, highlighted in green) (25). However, to our knowledge no other residues of the NAC domain have been shown experimentally to be required for protein dimerization, nor have the above residues been shown to have a biologically significant role *in planta*.

**Figure 1:**
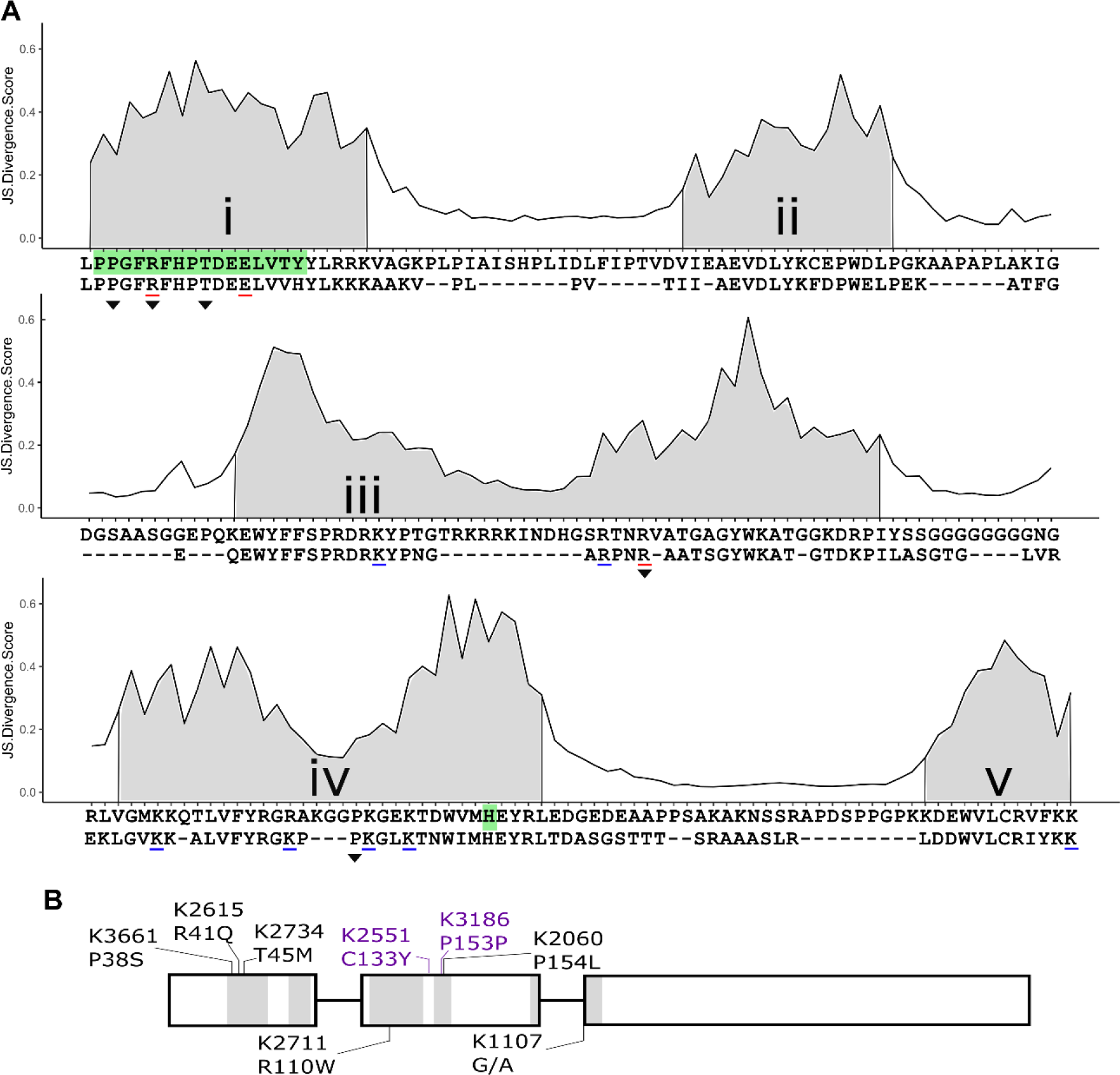
Identification of highly conserved residues of NAM-A1 in the cultivar Kronos TILLING population. The Jensen-Shannon Divergence scores for a consensus alignment of NAC-domain-containing proteins from seven plant species (see methods) highlight the conserved nature of the 5 NAC subdomains, highlighted in grey (A, i-v). Residues present in fewer than 10% of sequences across the set of NAC transcription factors were excluded from the alignment. Previous work has tested the role of specific residues in protein binding (subdomain i) and DNA binding (subdomains iii to v); residues which were required for the corresponding process are underlined in red, those that were not required are underlined in blue (24). Kang *et al.* showed that a histidine residue in subdomain iv plays a role in protein dimer stabilisation through interactions with part of the protein dimerization domain (subdomain i; both highlighted in green) (25). The consensus NAC domain sequence obtained from this alignment is shown below the conservation scores (first line). The aligned NAC-domain of *NAM-A1* is shown below the consensus sequence. Residues with missense mutations identified in the Kronos TILLING population are highlighted with a triangle (black). Note that the missense mutation in a non-conserved residue (C133Y) in K2551 is not present in the consensus alignment as it is not conserved across NAC proteins. Missense mutations, as well as non-conserved and synonymous controls (purple) and the splice acceptor variant K1107, are shown on a schematic of the *NAM-A1* structure (B).

In wheat, a NAC transcription factor (encoded by *NAM-B1*) was previously shown to act as a positive regulator of senescence (5). In modern-day hexaploid (bread) and tetraploid (pasta) wheat, this gene is either non-functional or deleted (5). Subsequent work in hexaploid wheat showed that a truncation mutant of the A genome homoeolog, *NAM-A1*, is sufficient in itself to cause a significant delay in leaf senescence (26). The loss of NAM-A1 also led to reduced grain protein content (GPC) in some of the environments tested, though this was less severe than in the D-genome truncation or the double A and D genome mutants. A similar truncation variant of *NAM-A1* was also shown to significantly delay leaf senescence in tetraploid wheat, and this also corresponded to a significant decrease in GPC (27). It is not clear, however, to what extent *NAM-A1* allelic variation in missense mutations may affect these phenotypes. Additionally, although these studies report that peduncles often remain green in *NAM-1* mutants, none have specifically explored the role of *NAM-A1* (or its homoeologs) in regulating senescence of the peduncle.

Recently, an *in silico* TILLING (Targeting Induced Local Lesions in Genome) resource has provided an unmatched source of point-mutation variation in both tetraploid and hexaploid wheat (28). Due to the polyploid nature of wheat, a higher dose of the mutagenizing chemical, ethyl methane sulphonate, could be used in the production of the TILLING population than typically used for diploid plants. This led to a mutation-dense set of lines that have been sequenced after exome-capture and are now easily interrogable through EnsemblPlants (29). When mapped against the latest genome reference (30), 99% of sequenced wheat genes have at least one missense mutation, with an average of 30 missense alleles per gene. This provides a useful resource for *in planta* studies of the roles of specific residues in protein function.

Here, we have leveraged the tetraploid *in silico* TILLING resource to study the impact of specific missense mutations on *NAM-A1*. Unlike most wheat genes which exist as functional copies on each of the two genomes in tetraploid wheat, the B-genome copy of *NAM-A1* is non-functional. As a result, we can study the impact of specific mutations on *NAM-A1* in a diploid context. We identified a set of five missense mutations in the NAC domain which are predicted to be highly deleterious (20, 24, 25). Based on these mutations, we characterised their impact on flag leaf and peduncle senescence and cell death *in planta*. We also assessed their ability to prevent protein binding *in vitro*. We identify four residues that lead to a delay in plant senescence and a loss of protein binding ability. We suggest that the function of these residues is likely conserved across species and that these residues are required for *NAM-A1* function.

## Results

### The TILLING population contains missense mutations in highly conserved residues of the *NAM-A1* NAC domain

To visualise the conservation of the NAC domain amongst the plant kingdom, we obtained the NAC domain sequence from 1320 NAC-domain-containing proteins from seven species across the plant kingdom (*Triticum aestivum, Hordeum vulgare, Oryza sativa* var. *japonica, Physcomitrella patens, Populus trichocarpa*, *Zea* mays, and *Arabidopsis thaliana*; Additional File 1). We utilised the Jensen-Shannon divergence (JSD) method, as implemented in Capra and Singh (31), to characterise the estimated conservation level of each residue in the aligned NAC domain (Figure 1A). Across the entire aligned NAC domain, we obtained a mean JSD value of 0.21. We then associated our consensus NAC domain sequence with the previously described subdomains (20). Breaking the NAC domain down into its subdomains, we found that the mean JSD value for each of the five subdomains is higher than that of the entire domain, ranging from 0.24 in the more heterogeneous subdomain iii, to 0.39 in the highly conserved subdomain i (Figure 1A). The higher levels of conservation in the subdomains is also evident in the comparison of the consensus NAC domain sequence (upper sequence, Figure 1A) with the aligned NAC domain sequence from the wheat NAC transcription factor *NAM-A1* (bottom sequence, Figure 1A).

Based on the highly conserved residues identified above, we selected five missense mutations present in cultivar Kronos TILLING lines which were predicted to have a deleterious effect on the protein function (Figure 1B, Table 1, Supplementary Table 1, Additional File 2). Three of the mutations, present in Kronos TILLING lines K3661, K2615 and K2734 are located at the predicted protein dimerization interface in subdomain i. The remaining two mutations (in lines K2711 and K2060) were located at the predicted DNA binding interface in subdomains iii and iv. A further two lines were selected as controls, with one encoding a synonymous mutation of a highly conserved residue (K3186) and the other a missense mutation in a non-conserved residue (K2551) between subdomains iii and iv. Finally, one line, K1107, was identified with a splice-junction acceptor mutation that leads to a predicted 11-residue deletion encompassing most of subdomain v (Supplementary Figure 1, Additional File 2,). Two of these mutations had previously been shown in Arabidopsis to have a role in either protein dimerization (K2615, R41Q) and DNA binding (K2711, R110W) (Table 1) (24). In the remainder of the paper, the TILLING lines will be referred to with both the TILLING line and corresponding mutation (e.g. K2711_R110W_). Line K1107, containing a splice acceptor variant (SAV), will be referred to as K1107_SAV_.

**Table 1.** Selected TILLING mutations in *NAM-A1*

| Line | SNP | RefSeqv1<br>chr6 (bp) | Consequence | Sub<br>domain | Mutation | SIFT | Region (20) | Consequence in<br>ANAC019 |
| --- | --- | --- | --- | --- | --- | --- | --- | --- |
| Kronos3661 | C/T | 77,100,016 | Missense | i | P38S | 0 | Dimerisation | - |
| Kronos2615 | G/A | 77,100,006 | Missense | i | R41Q | 0 | Dimerisation | Protein Binding (24) |
| Kronos2734 | C/T | 77,099,994 | Missense | i | T45M | 0 | Dimerisation | - |
| Kronos2711 | C/T | 77,099,581 | Missense | iii | R110W | 0 | DNA interface | DNA Binding (21, 24) |
| Kronos2551 | G/A | 77,099,511 | Missense | - | C133Y | 0.78 |  | - |
| Kronos2060 | C/T | 77,099,448 | Missense | iv | P154L | 0 | DNA interface | - |
| Kronos3186 | G/A | 77,099,450 | Synonymous | iv | P153P | NA |  | - |
| Kronos1107 | G/A | 77,099,212 | Splice Acceptor<br>Variant | v | SAV | NA | DNA interface | - |

### TILLING mutations lead to delayed senescence

M_5_ generation plants of the selected TILLING lines were grown and scored for visual senescence of the flag leaf and peduncle in glasshouse conditions (Figure 2, Supplementary Figure 2, Additional File 2). Across two separate experiments, none of the mutant lines showed a consistent delay in flag leaf senescence relative to the wild-type siblings. Flag leaf senescence onset was significantly delayed in four of the lines in Experiment 1 (two-sample Wilcoxon test, p < 0.05, Supplementary Figure 2, Additional File 2), however this difference was not recapitulated when repeated in Experiment 2 (Figure 2A). Relative flag leaf chlorophyll content of the wild-type and mutant lines was also not significantly different throughout senescence in Experiment 2 (Supplementary Figure 3, Additional File 2).

**Figure 2:**
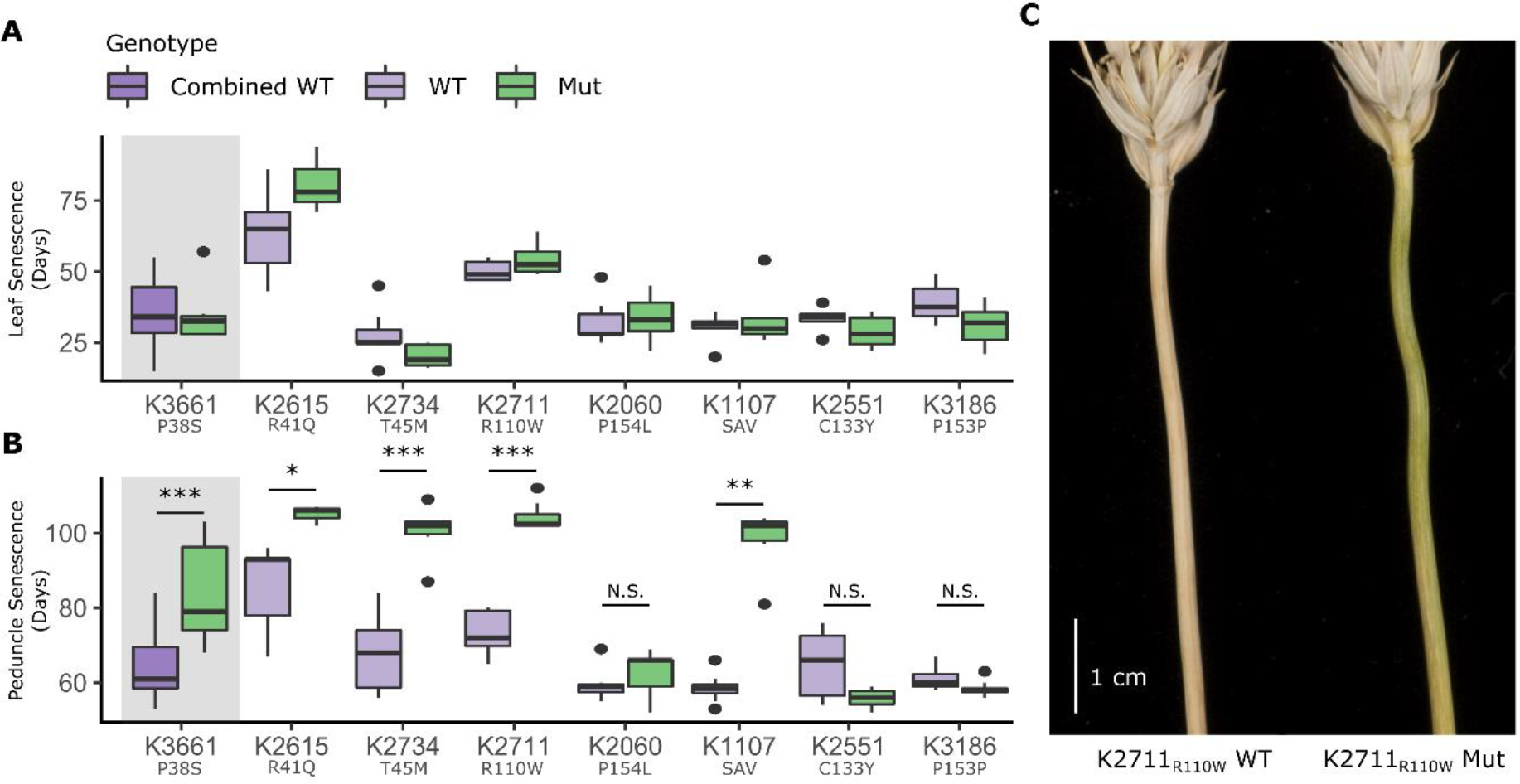
Mutations in the NAC domain of *NAM-A1* delay peduncle senescence in the glasshouse. TILLING lines were scored for flag leaf senescence (A) and peduncle senescence (B) under glasshouse conditions (Experiment 2). Note that K3661_P38S_, which was homozygous for the *NAM-A1* mutant allele, was compared to the combined data of all wild-type lines (see results for details; grey background). The Y-axis shows days to flag leaf (A) or peduncle (B) senescence after heading. The delay in peduncle senescence for K2711_R110W_ is pictured in (C) at 90 days post-anthesis. All statistics shown are two-sample Wilcoxon tests; *, p < 0.05; **, p < 0.01; ***, p < 0.001. N is between 8-10 plants for all lines except K2615 Mut where n = 3.

However, while flag leaf senescence was not consistently delayed in the *NAM-A1* mutants, this was not the case for peduncle senescence. Four lines, including three missense, K2615_R41Q_, K2734_T45M_, K2711_R110W_, and the splice mutation, K1107_SAV_, showed a highly significant delay in peduncle senescence in the mutant plants compared to the wild-type plants (Figure 2B and C, Supplementary Figure 2). In Experiment 2, K2711_R110W_ mutant plants took 104 days, on average, for peduncle senescence to occur following heading, compared to 73 days for the wild-type plants, a significant difference of 41 days. Similarly, peduncle senescence was significantly delayed by approximately 23, 33 and 40 days in the K2615_R41Q_, K2734_T45M_, and K1107_SAV_ mutants, respectively (all p < 0.05). K2060_P154L_ showed a more subtle delay in peduncle senescence, though this was only significant in Experiment 1 (p < 0.05, 4 days). A fourth line, K3661_P38S_, contained a homozygous mutation in the M_5_ plants, and as a result could not be compared directly to a wild-type line in the same mutant background. However, when compared to the data for all wild-type *NAM-A1* lines across the various alleles, K3661_P38S_ was significantly delayed in peduncle senescence, occurring on average 83 days after heading (p < 0.001). The synonymous mutation in K3186_P153P_ and non-conserved missense mutation in K2551_C133Y_ did not significantly delay peduncle senescence with respect to their wild-type siblings in either experiment.

Following the glasshouse experiments, we selected lines K2711_R110W_ and K1107_SAV_ for testing in field conditions. Across three years (2016, 2018 in the United Kingdom and 2017 in California, USA), K2711_R110W_ consistently showed a significant delay in both flag leaf and peduncle senescence (Figure 3A and B, Student’s t-test, p < 0.001). This effect was stronger than that seen for K1107_SAV_, which showed significant delays in peduncle senescence in all years (p < 0.001), but only in flag leaf senescence in 2018 (p < 0.05). Peduncle chlorophyll content was quantified in the 2018 field trial at 33 and 49 days after anthesis (DAA) (Figure 3C). At 33 DAA, chlorophyll levels were significantly higher in the K2711_R110W_ mutant plants than in wild-type (Student’s t-test; p < 0.05). By 49 DAA, the levels of chlorophyll A in both the wild-type lines had dropped to zero, while chlorophyll was significantly higher in both *NAM-A1* mutant lines (K2711_R110W_ and K1107_SAV_, p < 0.05).

**Figure 3:**
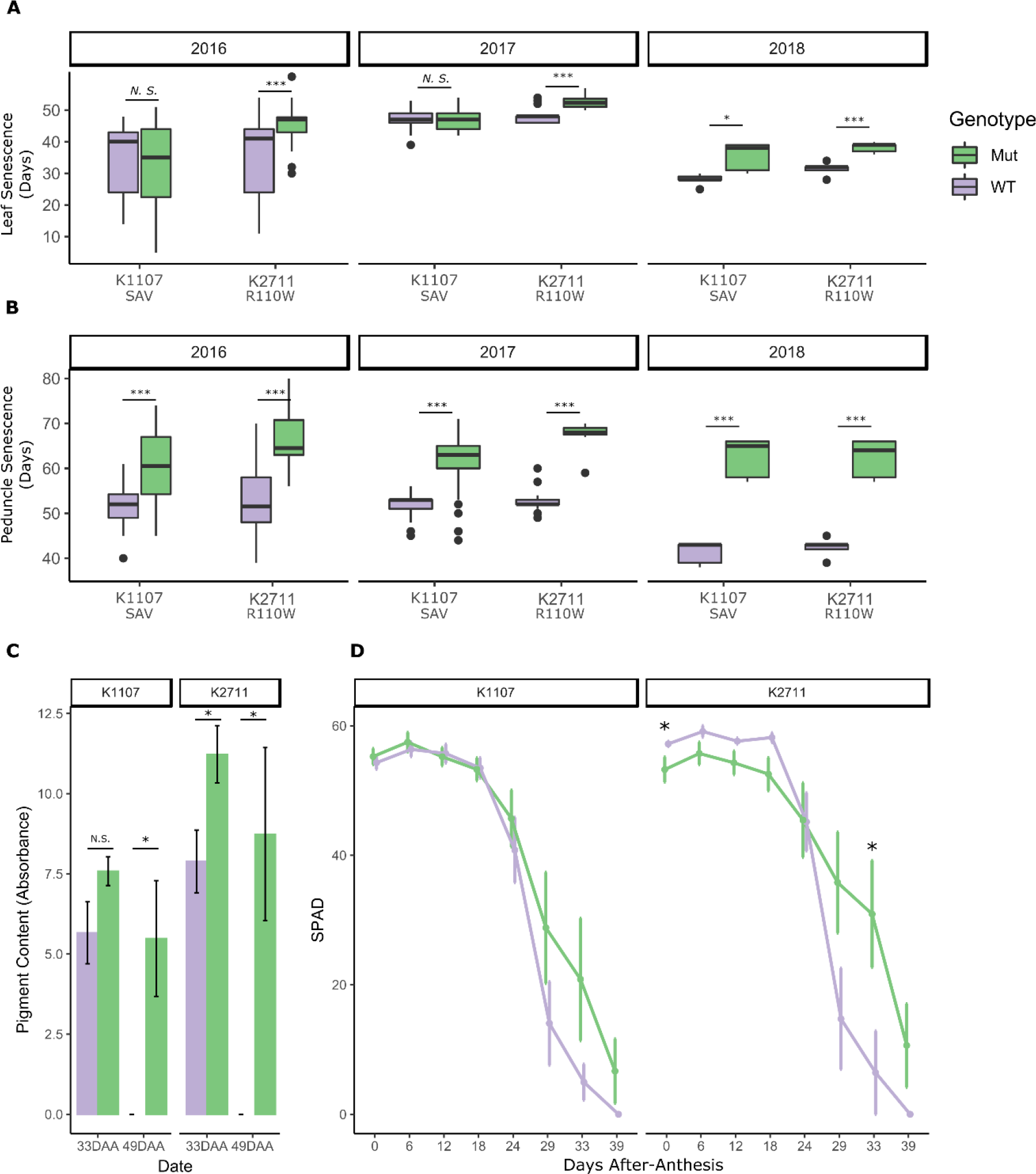
Mutations in NAM-A1 lead to a significant delay in peduncle senescence in the field. TILLING lines K1107 and K2711 were scored for flag leaf (A) and peduncle (B) senescence in the field across three years. The Y-axis shows days to flag leaf (A) or peduncle (B) senescence after heading. Peduncle chlorophyll content (C) was quantified at 33 and 49 days after anthesis (DAA) for the wild-type and mutant plants of each line in the 2018 field trial. SPAD measurements of flag leaf chlorophyll content (D) were carried out on the lines in 2018 throughout senescence. See methods for sample size each year. Statistical comparisons were carried out with a Student’s t-test for all panels except the SPAD time course, which used the Kruskal-Wallis Rank Sum test; *, p < 0.05; **, p < 0.01; ***, p < 0.001.

Flag leaf senescence progression was also quantified in the 2018 field trial using SPAD measurements. From anthesis to approximately 24 DAA, no significant difference was observed between the mutant or wild-type plants; though K2711_R110W_ wild-type lines contained significantly more SPAD units at anthesis, this difference was not maintained in the following timepoints (Figure 3D). At 29 DAA, separation could be seen between the mutant and wild-type plants for both mutant lines. However, only K2711_R110W_ showed a significant difference in flag leaf SPAD units between the mutant and the wild-type, at 33 DAA (Kruskal-Wallis Rank Sum test, p < 0.05). This is consistent with the stronger flag leaf senescence phenotype observed in K2711_R110W_ compared to K1107_SAV_.

### TILLING mutations lead to reduced protein content and no change in grain size parameters

Grain samples were taken from the K1107_SAV_ and K2711_R110W_ lines grown in 2016 and 2018 (UK), and grain size parameters were measured. No consistent effect on grain size parameters was observed in the field for either K2711_R110W_ or K1107_SAV_ in either of the two years (Supplementary Figure 4, Additional File 2). This lack of grain size effect is consistent with that observed previously in hexaploid wheat for a NAM-A1 truncation allele (26).

**Figure 4:**
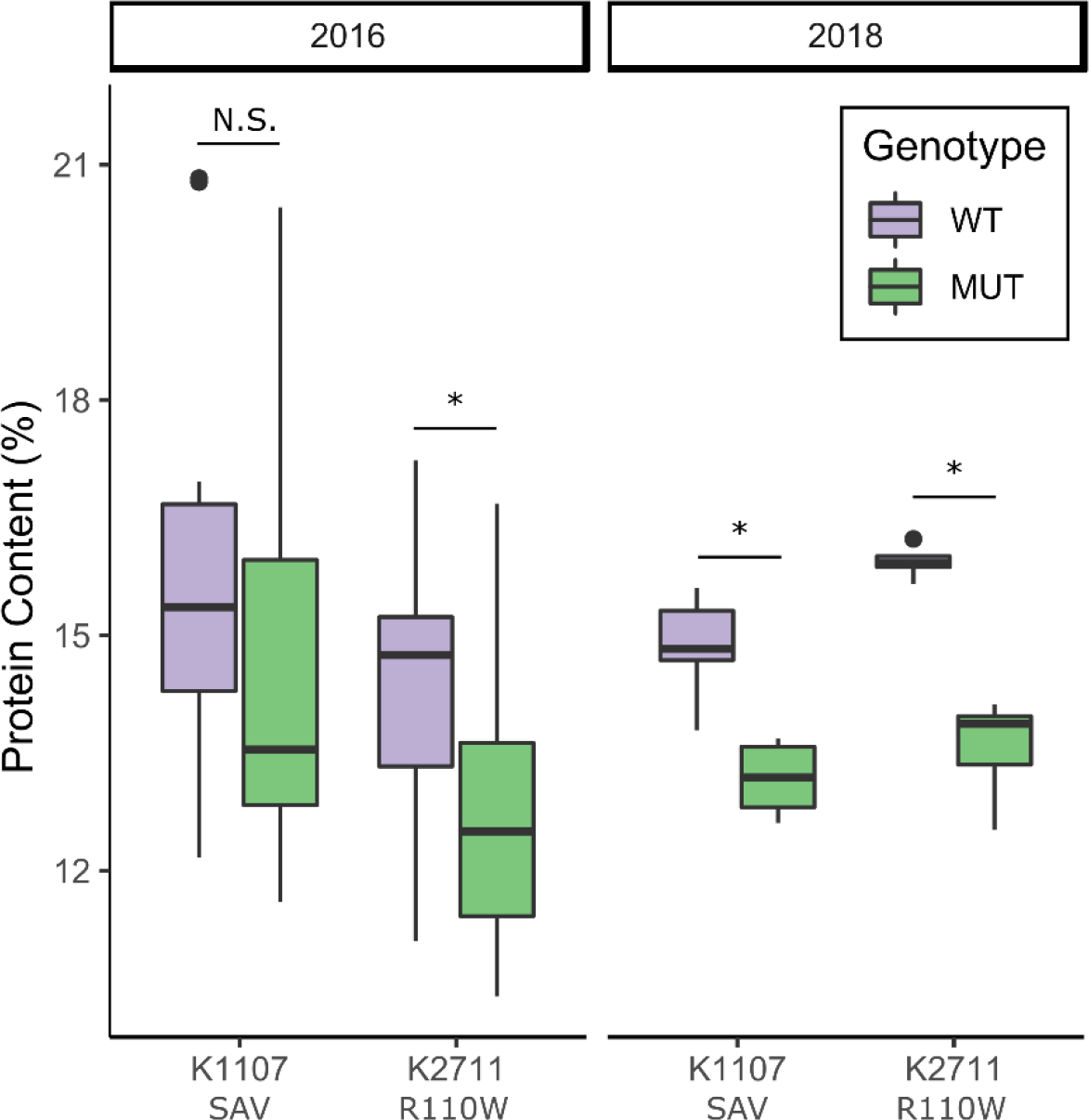
Grain protein content is decreased in the K2711_R110W_ mutant line under field conditions. Grain protein content was measured from mature grain samples taken from the 2016 and 2018 field trials of lines K1107_SAV_ and K2711_R110W_. Statistical comparisons were carried out with the Wilcox Test; *, p < 0.05. N = 5 for 2018 field trials and varies between 10 and 22 for the 2016 field trials.

Near-infrared (NIR) measurements of grain protein were carried out on the same set of grain samples from field trials of K2711_R110W_ and K1107_SAV_. In 2016, the mutant plants from the K2711_R110W_ population showed a significant decrease in grain protein content (GPC) compared to wild-type plants (p < 0.05, Figure 4). The K1107_SAV_ mutant plants, however, showed no consistent reduction in GPC. In 2018, the reduction in GPC in the mutant lines was consistent across the K2711_R110W_ and K1107_SAV_ populations (14.7% and 11.5% GPC reduction, respectively, p < 0.05, Figure 4). These results are consistent with previous work in hexaploid wheat, where truncation mutants in *NAM-A1* and *NAM-D1* led to reductions in GPC content in the field, although this varied between years (26).

### Mutations in the protein-binding domain prevent heterodimerisation

To determine how the missense mutations were causing a delay in senescence, we characterised their impact on protein function. Previously, a yeast two-hybrid (Y2H) screen had been carried out on the wheat NAM-B1 protein which identified the NAM-A1 homoeolog as a putative binding partner (unpublished results). We confirmed this interaction via yeast two-hybrid and co-immunoprecipitation, demonstrating that NAM-B1 and NAM-A1 can interact *in vivo* (Figure 5, Supplementary Figure 5, Additional File 2). We then used this known interaction to test whether the selected mutations in the conserved residues of NAM-A1 (Table 1) resulted in a loss of protein interaction to NAM-B1.

**Figure 5:**
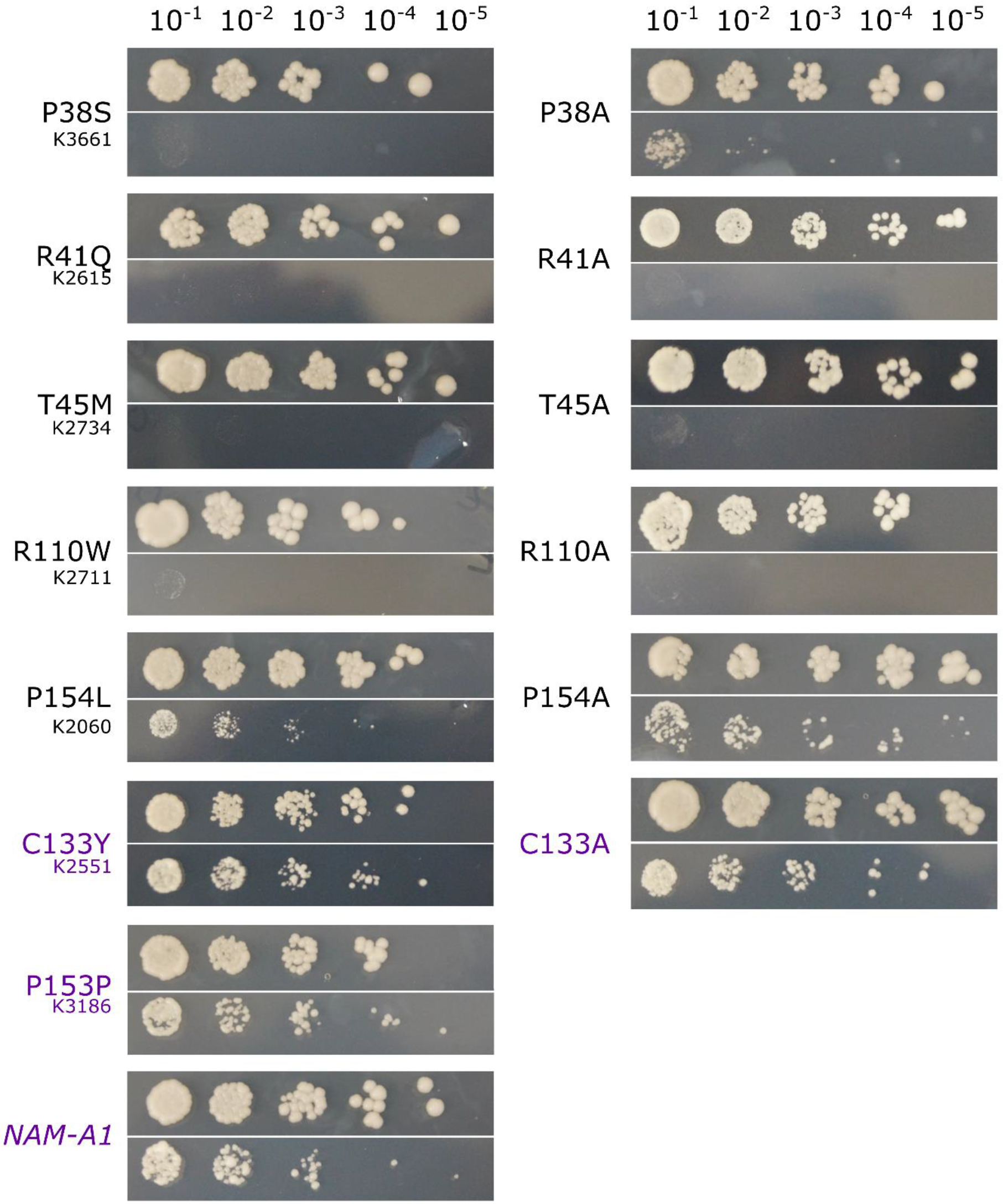
Mutations in the protein-binding domain of NAM-A1 inhibit protein binding. Yeast-two-hybrid interaction screens between NAM-A1 alleles and wild-type NAM-B1 were tested on control (SC-LT; top row) and selective (SC-LTH + 10mM 3AT; bottom row) media at ten-fold OD600 dilutions, from 10^−1^ to 10^−5^. Growth on selective media indicates interaction between the NAM-A1 allele and wild-type NAM-B1. Lack of growth on selective media indicates a loss of interaction. All controls showed equivalent growth on selective media as the wild-type *NAM-A1* (purple). Left hand panels correspond to TILLING mutations whereas right hand panels are alanine substitutions. The corresponding TILLING line is shown underneath the mutation on the left panel.

Mutant alleles of NAM-A1 were cloned with the corresponding mutation found in the Kronos TILLING lines, as detailed in Table 1. The same residues were also mutated to an alanine residue to test whether any phenotype was due to the specific mutation in the TILLING line, or due to functional properties of the conserved residues themselves (32). Only the NAC domains of the NAM-A1 alleles were used in the Y2H to prevent auto-activation of the yeast GAL4 promoter by the C-terminal transcriptional activation domain (Additional File 3)

Screening of the interactions on selective media found that of the five missense mutations in conserved residues, four led to a complete loss of interaction with both the TILLING and alanine mutation (Figure 5, Supplementary Figure 6, Additional File 2). This included the three mutant alleles in the predicted protein-binding domain (P38S, R41Q, T45M) and one of the two alleles in the predicted DNA-binding domain (R110W). One mutation (P154L) only partially reduced the ability to bind NAM-B1 as it showed a reduced level of interaction for the TILLING mutant compared to the controls. However, this interaction was fully recovered for the alanine mutation. All controls retained interaction in the selective conditions, including the non-conserved missense mutation C133Y in K2551.

### Missense mutations reduce cell-death induction

Following the Y2H screen, we transiently over-expressed the *NAM-A1* alleles in *Nicotiana benthamiana* to determine their effect on the cell death response (see methods). Cell death was scored on both chlorosis (under white light) and the build-up of phenolic compounds leading to auto-fluorescence (under UV light) (33). Over-expression of wild-type *NAM-A1* leads to a cell death phenotype similar to that observed during a weak hypersensitive response, with evidence of chlorosis and auto-fluorescence significantly higher than the mock (p < 0.001, Figure 6). This is consistent with the role of *NAM-A1* in wheat as a positive regulator of senescence (5, 26, 27).

**Figure 6:**
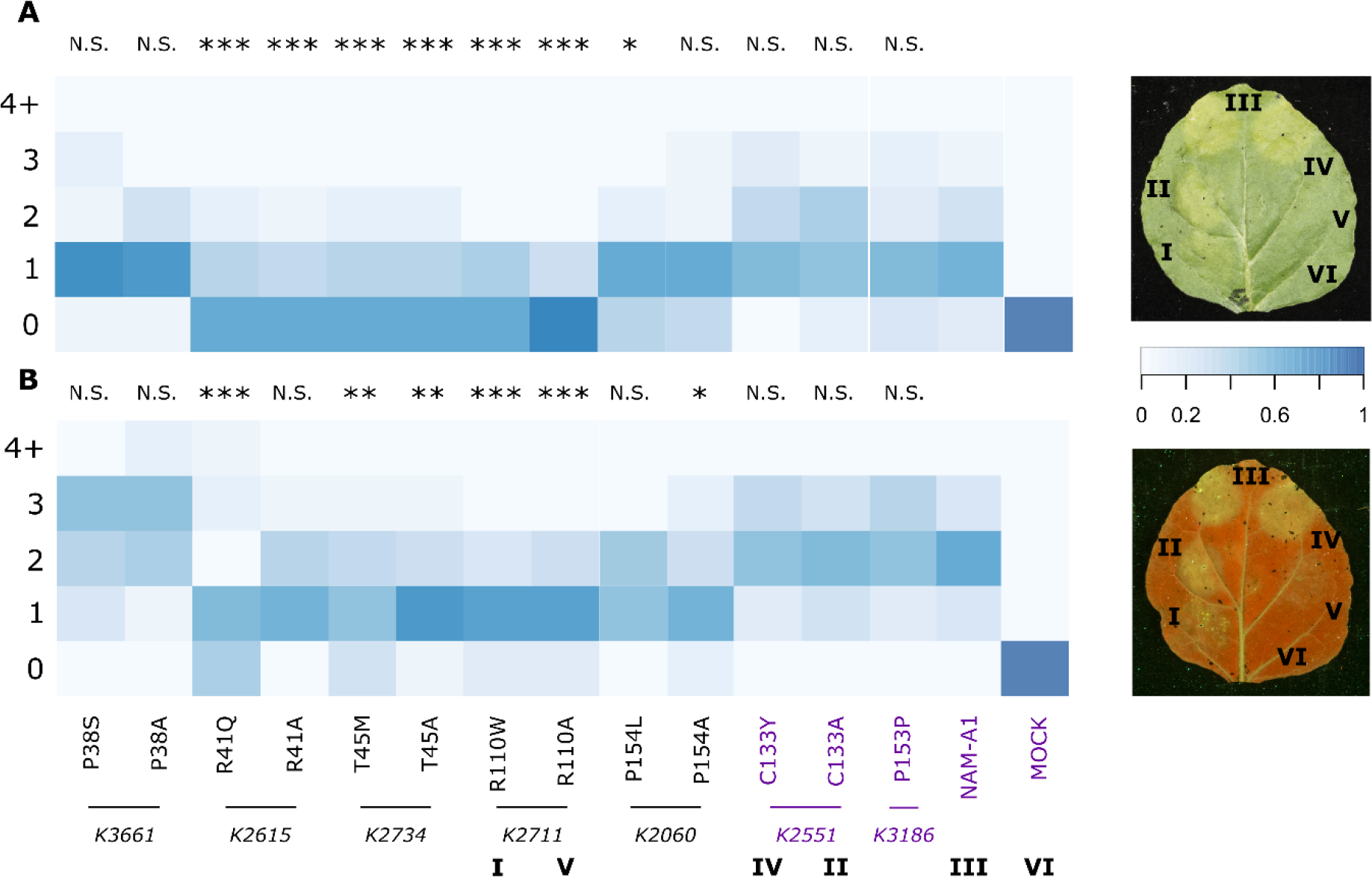
Mutant alleles of *NAM-A1* show reduced cell death response. Heterologous expression of the mutant *NAM-A1* alleles in *N. benthamiana* shows a significant reduction in cell death response for both chlorosis (A) and cellular breakdown (B) compared to the wild-type allele. Cell death for chlorosis and necrosis are scored on a scale derived from Maqbool *et al.* (33), whereby 0 is no chlorosis or necrosis and 6 is a high level of chlorosis or necrosis. All control alleles (purple) do not differ significantly from the wild-type. The corresponding TILLING line is shown below the TILLING/alanine mutant pairs. A typical image of a patch-infiltrated leaf is shown for both chlorosis scoring (top, adaxial) and UV-scoring (bottom, abaxial); number labels correspond to those beneath the heatmap; M is mock treated. The UV image (bottom) has been flipped horizontally to allow comparison with the white-light image. Heatmap scale is the fraction of total replicates which fell into each of the 5 bins. Note that all scores greater than or equal to 4 were assigned into the same bin (“4+”) due to low incidence; n varies from 20-35. Statistical comparisons shown are between the mutant allele and wild-type *NAM-A1* based on a linear model fit to the Poisson distribution. *, p < 0.05; **, p < 0.01; ***, p < 0.001.

We then screened the *NAM-A1* mutant alleles, including TILLING and alanine mutants, for reductions in the cell-death response. In general, we found that both methods of scoring cell death (chlorosis and auto-fluorescence) corresponded closely with the loss of protein interaction observed in the Y2H screen. Of the four mutant alleles that showed a complete loss in protein interaction in yeast (Figure 5), three also showed significantly attenuated chlorosis and cell death response compared to wild-type NAM-A1 (Figure 6, p < 0.01; R41Q, T45M, R110W). Only P38S, which showed loss of protein interaction, had no effect on cell death response. The P154L substitution, which partially reduced protein interaction, was also borderline significant (p = 0.03) in its chlorosis response. Both control alleles show no decrease in either chlorosis or cell death, including the non-conserved missense mutation (C133Y). Protein expression of all constructs was verified using a Western blot (Supplementary Figure 7, Additional File 2)

## Discussion

NAC transcription factors have been characterised across several plant species (22, 23, 34) and found to have central roles regulating a variety of stress and developmental responses, including senescence. In this study, we have identified and characterised a set of residues which are required for protein interaction between the wheat NAC transcription factor NAM-A1 and its homoeolog NAM-B1.

### Highly conserved residues of the NAC domain are required for protein function across species

While the NAC domain is conserved across species, it is yet to be determined if the same residues are required for protein function between species and indeed across NAC TFs in the same species. Here we characterised the effect of missense mutations in conserved residues on protein function. Olsen *et al.* (2005) previously showed that a mutation in a highly conserved salt-bridge-forming arginine residue in Arabidopsis NAC transcription factor ANAC092 (R19) is sufficient to disrupt protein dimerization (24). Here we have independently confirmed the importance of this residue (R41 in NAM-A1) in protein-protein interaction (R41Q; Figure 5) and show that the mutation in this residue leads to a delay in wheat senescence (K2615; Figure 2) and reduced cell death response in *N. benthamiana* (Figure 6). Similarly, the R110W mutation in a second arginine in subdomain iii is located at the same residue as that identified in ANAC019 (R88) to be essential for DNA binding (24). Here we show that this residue is also important for protein-protein interactions, a somewhat unexpected result given the residue’s location in the putative DNA-binding domain of the protein. Furthermore, we characterised the effect of this residue *in planta* and show that the R110W substitution in K2711 leads to a significant delay in wheat flag leaf and peduncle senescence in the field, as well as a reduction in cell death response in *N. benthamiana*.

During the course of this study, Kang *et al.* investigated the role of a specific residue in stabilising protein structure at varying pH levels in the Arabidopsis protein ANAC019 (25). These authors report that the protonation status of the H135 histidine residue of ANAC019 (Figure 1A; highlighted green in subdomain iv) defines the dimer form of the protein and its ability to form a complex with DNA. Based on protein structural simulations, the authors suggest that a “perfect” dimer is formed when the H135 residue is not protonated, preventing formation of a salt-bridge with the D24 residue in subdomain i. In this state, various chemical interactions occur between residues of the dimerising proteins which help stabilise the “perfect” dimer (Figure 1A). If the H135 residues is protonated, it is predicted to form the salt-bridge with residue D24 which disrupts the majority of these interactions. Given these results, it seemed plausible that the residues highlighted as forming intra-dimer bonds in the “perfect” dimer state may be essential for protein dimerization.

Three of the mutations we studied here are located within the dimerization domain highlighted in Kang *et al.* (Figure 1A; green highlight in subdomain i)(25). Two residues, P38S (K3661) and R41Q (K2615), correspond directly to residues predicted to form stabilising bonds in the “perfect” dimer state (P16 and R19 in ANAC019, respectively). The third mutation, T45M (K2734), falls within this dimerization region and is located adjacent to the critical salt-bridge-forming residue (D24) in the NAC consensus sequence (Figure 1A). In this study we find that mutations in these three residues of NAM-A1 lead to complete loss of protein interaction ability and delayed peduncle senescence. These results are consistent with the predicted role of these residues in stabilising the formation of the NAC dimers found in Kang *et al.* and provides strong evidence for their biological relevance *in planta* (25). Our ability to both recapitulate and support these findings in wheat also highlights the fact that the functional domains, and essential residues, of NAC transcription factors are likely to be highly conserved across species.

### The Kronos TILLING population facilitates studies of novel allelic variation in wheat

In this study, we have also shown that the wheat TILLING population, particularly the tetraploid Kronos population, provides an invaluable resource to study large allelic series of mutations in wheat. Using *NAM-A1* facilitated this study as it functions as a single-copy gene in tetraploid wheat (given that the homoeolog *NAM-B1* contains a frame-shift leading to non-functional protein in Kronos) (5, 27). As a result, we were able to characterise the M_5_ mutant lines directly without confounding effects due to homoeolog redundancy. Approximately 60% of genes with at least one mutation contain at least one premature truncation mutant in the Kronos population (28). This suggests that for a large portion of genes, it should be possible to study a similar allelic series through single crosses between desired missense mutations in one homoeolog, and a common premature truncation mutant in the other homoeolog.

Carrying out studies of allelic series such as this one in wheat, and indeed in other crops, can help identify specific alleles that lead to a gradient of phenotypes. We found that the effect of the mutations varied from a few days delay to over 20 days delay in peduncle senescence in the glasshouse. Similarly, the strength of the phenotype observed between K2711_R110W_ and K1107_SAV_ in the field also varied, particularly for flag leaf senescence. We are currently developing germplasm to test the effect of the additional missense mutations on senescence in field trials. It is known that allelic variation within populations can account for variable success across environments (35–37). Indeed, previous studies of natural variants in *NAM-A1* identified a missense mutation (haplotype NAM-A1c) which was associated with an intermediate grain protein content phenotype between the wild-type allele and a third haplotype containing the missense mutation in tandem with a frame-shift mutation in the C-terminal domain (38). This missense mutation sits just beyond the end of subdomain iii, in a relatively un-conserved residue. As additional agronomically-important genes are identified, it will become increasingly possible to mine the TILLING population for novel variation, or base-edit alleles (39), to intelligently introduce variation based on predicted consequences in conserved domains.

### NAM-A1 mutants have a more severe effect on peduncle senescence than on flag leaf senescence

Our studies with the allelic series of *NAM-A1* mutants have also highlighted that while mutations in *NAM-A1* can lead to delays in flag leaf senescence, particularly in the field, there is also a strong and significant effect on peduncle senescence both in glasshouse and field trials. It is worth noting that the traits scored for visual senescence in flag leaves and peduncles do differ. Flag leaf senescence is scored when 25% of the leaf has turned yellow—a measure of senescence onset. In contrast, peduncle senescence is scored when the top inch of the peduncle is fully yellow—a measure of senescence termination. This difference is due to the difficulty inherent in determining when a peduncle begins to senesce. Most often senescence in the flag leaf occurs in a directional manner, moving from the tip of the leaf towards the base. Peduncle senescence, in contrast, can occur around the stem simultaneously, making it difficult to assign a clear onset location and timing for senescence. However, the measure of flag leaf senescence progression via SPAD measurements allows senescence termination to also be assessed for the flag leaf. This enables a more meaningful comparison with the visual measure of peduncle senescence. Significant delays in peduncle senescence were seen in the glasshouse experiments with the *NAM-A1* mutants (Figure 2B). However, no significant delays in flag leaf senescence progression were observed for any of the mutants using SPAD measurements (Supplementary Figure 3, Additional File 2). Similar results were also observed for the *NAM-A1* mutants in the field (Figure 3B and 3D). This data corroborates that observed when comparing the visual senescence scores; namely that the *NAM-A1* mutants have a more severe and substantial effect on peduncle senescence than on flag leaf senescence in both glasshouse and field experiments.

Previous studies on the *NAM* genes have predominantly focused on the flag leaf senescence phenotype, though the delayed peduncle senescence phenotype has been noted (5, 26, 27, 40). It is known that the *NAM* genes regulate the remobilisation of nutrients into the spike, which must pass through the peduncle before reaching the grain. Peduncle nitrogen content at harvest is higher in *NAM-1* knock-out lines, while NAM RNAi knock-down lines show altered profiles of micronutrient levels in the grain (26, 41). Earlier work failed to identify any expression of the *NAM* genes in peduncle tissue at 14 DAA, initially indicating that the NAM genes might be acting indirectly on nutrient remobilisation and senescence in the peduncle (42). However, more recent RNA-Seq data from a comprehensive developmental time course has identified expression of the *NAM* genes in the peduncle during grain filling (Supplementary Table 2, Additional File 2) (43, 44). At the latest timepoint available for peduncle tissue (milk grain stage, approximately 15 DAA), the levels of all *NAM* genes are higher in the peduncle than they are in the flag leaf, supporting a direct role for the *NAM* genes in the peduncle. This corroborates more recent expression analysis which found higher expression levels of the *NAM* genes in the peduncle compared to the flag leaf blade from 7 DAA (45). It’s tempting to speculate that by delaying senescence processes in the peduncle, the *NAM* genes are hindering the transport of nutrients into the grain, instead sequestering them in the peduncle. Support for this hypothesis comes from work on RNAi knockdown lines of the *NAM* family that has shown that the mutant lines retain substantially more iron, zinc, nitrogen, and fructan content in the peduncle during senescence than wildtype lines (26, 41, 46).

The focus on understanding flag leaf senescence compared to that of other tissues such as the peduncle is understandable given that the flag leaf provides the majority of sugars and nutrients remobilised to the grain (47, 48). Additionally, the progression of flag leaf senescence is relatively easy to measure using the non-destructive SPAD meter. Whereas, beyond visual scoring of yellowing, quantitative scoring of peduncle senescence relies on destructive measurements of pigment content. However, our results suggest that understanding the mechanisms governing the timing and regulation of peduncle senescence is also necessary to understand how flag leaf senescence impacts on grain quality and yield.

### Conclusions

Here we have combined *in silico* prediction of critical residues in the NAC domain with biochemical and *in planta* studies of their role in the function of *NAM-A1*. Using the Kronos TILLING population, we were able to rapidly identify and screen missense mutations of *NAM-A1* in the greenhouse and in the field. After identifying four mutants that caused delayed peduncle senescence, we used the yeast 2-hybrid system to show that the mutations lead to a loss of protein interaction and suggest that this is the mechanism by which the mutants disrupt protein function. The characterisation of these mutations confirms previous work on NAC TFs carried out in Arabidopsis and identifies previously uncharacterised residues which are required for protein function. Together, these results highlight our ability to use new genetic resources in wheat, such as the TILLING population, to interrogate the effects of specific mutations on gene function and phenotype. Moving forwards, this presents a method for in-depth characterisation of other conserved protein domains using wheat which can then be integrated with knowledge generated in other species.

## Methods

### Plant Growth and Phenotyping

#### Glasshouse Experiments

Kronos TILLING mutant lines (49) containing mutations in the *NAM-A1* gene were selected as described previously (28). Briefly, lines containing missense mutations predicted or known to affect protein function were identified, alongside control lines with synonymous, non-conserved missense, or splice-variant mutations (Table 1) (20, 24). M_5_ seeds of the selected TILLING lines were pre-germinated on moist filter paper for 48h at 4°C. Seeds were then sown into P96 trays in 85% fine peat with 15% horticultural grit. At the 2-3 leaf stage, individual plants were transplanted to 1L round pots in Petersfield Cereal Mix (Petersfield, Leicester, UK). Plants were grown in standard glasshouse conditions, with 16:8 hours of light:dark cycles, and genotyped to identify homozygous mutant and wildtype lines (see genotyping methods below). The primary spikes were tagged at heading (75% spike emerged from flag leaf sheath; Zadoks growth stage 57) (50). Flag leaf senescence onset was scored when 25% of the main flag leaf had lost chlorophyll and appeared visually yellow. Peduncle senescence was scored when the top inch of the main peduncle became fully yellow. Experiments were repeated twice under the same conditions (referred to as Experiment 1 and 2 in the text).

#### Nicotiana benthamiana Growth

Seedlings were sown into half seed trays in F2 soil (Peat with 2.5kg/m^3^ Dolimite, 1.3kg/m^3^ Base fertiliser, 2.7kg/m^3^ Osmcote 3-4 months, 0.25kg/m^3^ Wetting agent, and 0.3kg/m^3^ Exemptor). Plants were transplanted 11 days after sowing into FP8 pots. Plants were grown under a 16:8 light:dark regime under sodium lamps (400W), with temperature set to 25°C day and 22°C night.

#### JIC Field Trials

Two lines (K2711 and K1107) were selected for further evaluation in the field. In each case, homozygous *NAM-A1* mutant plants were crossed to the wild-type original parent, Kronos. F_1_ plants were self-pollinated to produce F_2_ populations segregating for the mutant or wild-type *NAM-A1* allele. Two independent populations were obtained for K1107 (referred to as K1107-4 and K1107-6) and one population was obtained for K2711. The F_2_ populations were sown at Church Farm, Bawburgh (52°38′N 1°10′E), in March 2016. A total of 119 kg/ha of Nitrogen was applied during the 2016 growing season.

To score segregating F_2_ plants for phenotype and genotype, individual F_2_ seeds were hand-sown in 6×6 1 m^2^ grids of 36 individual plants with approximately 17 cm between each plant (K2711, n = 158 F_2_ plants; K1107-4, n = 218; K1107-6, n = 212). The individual F_2_ plants were scored for heading date (two tillers with spikes 75% emerged), leaf senescence (two tillers with flag leaves 25% senesced from tip), and peduncle senescence (two peduncles, top inch senesced). Senescence scoring was consistent with that used in the glasshouse experiments. Following identification of homozygous mutant and wild-type F_2_ individuals (see genotyping methods below), we selected between 13-16 individual plants for each line and genotype to take forward for further phenotyping in 2017 (Davis). A subset of these selected homozygous lines was also used for phenotyping in 2018 (JIC). Only individuals genotyped as homozygous wild-type or mutant were included in data analysis for 2016.

In April 2018, homozygous lines derived from independent F_2_ plants in the 2016 field trial were sown at Church Farm for K2711 (WT *NAM-A1*, n = 5; Mutant *NAM-A1*, n = 5) and K1107 (WT *NAM-A1*, n = 5; Mutant *NAM-A1*, n = 5). Each of the lines was considered an independent biological replicate of the mutant or wild-type genotype to account for the potentially confounding influence of segregating background mutations derived from the original TILLING mutants. Each biological replicate was sown in double 1 m rows, separated by a single empty row, arranged in a complete randomized design. Whole rows were scored for senescence using a similar method as detailed previously, however here we scored the double rows as “senesced” when 75% of the main tillers exhibited the leaf or peduncle senescence phenotype. A total of 124.5 kg/ha of Nitrogen was applied to the 2018 field trials.

#### Davis Field Trial

Homozygous lines derived from individual F_2_ plants were sown for K2711 (WT *NAM-A1* n = 13, Mutant *NAM-A1* n = 15) and K1107 (WT *NAM-A1* n = 14, Mutant *NAM-A1* n = 16) in November 2016 at the University of California Field Station, Davis, California (38° 31′ N, 121° 46′ W). As above, each of the lines was considered an independent biological replicate of the mutant or wild-type genotype. K2711 and K1107 were sown as two independent complete randomized design trials. Each biological replicate was sown as double 1 m rows separated by one empty row, identical to the JIC 2018 trials. Senescence was scored in the same manner as for the 2018 JIC trials, detailed above. A total of 200 lb N/acre were applied as ammonium sulphate, with half at pre-planting and the rest at the beginning of jointing (Zadoks 30, March 31 2017). Both the Davis and JIC trials were sprayed with appropriate commercial fungicides to avoid disease incidence.

#### Chlorophyll Content

Relative chlorophyll content was measured in leaves non-destructively using the SPAD-502 chlorophyll meter (Konica Minolta). For glasshouse trials, readings were taken eight times along the flag leaf blade and averaged to obtain an overall reading for each leaf (n = 5 for each line and genotype). For 2017 and 2018 field measurements, at least 2 individual leaves per biological replicate (n = 5 for each line and genotype) were measured as above and averaged for each time point.

Chlorophyll content was also measured chemically from sampled peduncle tissue in the JIC 2018 field trial. The top inch of peduncles was sampled at 33 and 49 days after anthesis and was cut into small segments (approximately 0.5 cm in length). Two peduncles were sampled per biological replicate and pooled before chlorophyll extraction. Five biological replicates were sampled for each homozygous genotype (K2711 WT, K2711 mutant, K1107 WT, and K1107 mutant). Plant tissue was soaked in 3 mL N,N-Dimethylformamide (analytical grade, Sigma Aldrich, UK) for 4-5 days at 4 °C until all pigment had leached from the tissue. Chlorophyll and carotenoid content were quantified as previously described (51).

#### Grain Size

Grain samples from individual plants (Glasshouse and Field 2016) and pooled samples (Field 2017 and 2018) were analysed using the MARVIN seed analyser (GTA Sensorik GmbH). Grain morphometric parameters, including area, length, and width, alongside approximate thousand grain weight (TGW) were obtained.

#### Grain Protein Content

Grain protein content measurements were taken using the Perten DA7250 NIR analyser. Grain samples of between 20 – 700 grains were tested for protein content (Protein dry basis %). Samples with fewer than 300 grains were analysed using a static cup; samples with more than 300 grains were analysed using a rotating cup.

#### Genotyping

To validate the presence of the *NAM-A1* TILLING mutations, we used KASP genotyping as previously described (52). Primers for genotyping were designed using Polymarker, where possible, or else designed based on aligned sequences of the *NAM-A1* homoeologs to ensure homoeolog specificity (Supplementary Table 3, Additional File 2) (53).

#### NAC domain alignment

NAC domain-containing proteins in *Arabidopsis thaliana*, *Zea mays*, *Populus trichocarpa*, and *Physcomitrella patens* were extracted from EnsemblPlants, using the Interpro ID IPR003441. NAC proteins from *Hordeum vulgare* and *Oryza sativa* were obtained from the curated list in Borrill *et al.* 2017 (22). NAC-domain containing proteins from *Triticum aestivum* were obtained from the gene annotations for RefSeqv1.1 (30).

Sequences were aligned using Clustal Omega v1.2.0, and manually curated in Jalview v2.10.5 (54). Following alignment, residues with less than 10% occupancy across the set of NAC transcription factors were excluded from further analysis, as in Borrill *et al.* 2017 (22). The non-conserved C-terminal domain was also excluded from further analysis. Based on this curated alignment, the Jensen-Shannon Divergence score at each residue was calculated as in Capra and Singh (2007), using the “Protein Residue Conservation Prediction” web server (31).

#### *NAM-A1* and *NAM-B1* Allele Cloning

TILLING mutations in the *NAM-A1* gene (TraesCS6A02G108300) were identified using the TILLING database at www.wheat-tilling.com (28). Specific alleles were selected based on conservation of the residue in the NAC domain (PSSM viewer, pfam02365) and predicted SIFT scores (55). For simplicity, the Kronos mutant lines are labelled as ‘K’ followed by their four-digit identifier (e.g. Kronos 2711 is referred to as K2711).

The *NAM-A1* sequence from *T. aestivum* (Transcript TraesCS6A02G108300.2) was synthesized into the pUC57 vector (Genewiz) and cloned into the pCR8 vector using the TOPO cloning kit using primers in Supplementary Table 4, Additional File 2. Site-directed mutagenesis was carried out using the primers in Supplementary Table 5, Additional File 2, to acquire the allelic series of *NAM-A1* variants. All cloned products were confirmed using Sanger sequencing (Eurofins Genomics). The *NAM-B1* sequence was obtained from *T. turgidum subsp. dicoccoides* (GenBank accession DQ869673.1) and was synthesized as detailed above (Additional File 3).

#### Co-Immunoprecipitation

The wild-type alleles of *NAM-A1* in the pGWB21 vector (N-terminal Myc tag) and *NAM-B1* in the pGWB12 vector were co-infiltrated into *N. benthamiana*, alongside P19 at a total OD of 0.6 and individual OD of 0.2 per construct. Leaves were harvested and snap-frozen at 3 days post-infiltration. Protein was extracted in a standard buffer (100mM Tris-HCl, pH 7.5; 150mM NaCl; 0.1% (v/v) Triton x-100; 1% (w/v) PVPP; 5mM EDTA; 10% glycerol; 2mM DTT; 1% protease inhibitors (Sigma)). 1 mL of protein extraction was applied to M5 anti-FLAG magnetic beads (Sigma). The beads were then boiled and the resulting fraction was run on a 12% SDS-PAGE gel, alongside the input and unbound fractions. The proteins were transferred to a nitrocellulose membrane and probed with both an anti-FLAG HRP conjugated antibody and an anti-Myc HRP conjugated antibody (Abcam). Total input was visualised by staining the membrane with Ponceau following antibody probing. The experiment was repeated three times with consistent results.

#### Yeast-Two-Hybrid

The ProQuest Two-Hybrid system was used with minor modifications to verify the presence or absence of protein interactions between *NAM-B1* and the *NAM-A1* alleles. Briefly, the sequences coding for the NAC domains of NAM-A1 and NAM-B1 (residues 1-217 and 1-215, respectively; Additional File 3) were amplified and cloned into pCR8, using the primers in Supplementary Table 4, Additional File 2. Alleles of *NAM-A1*(1-217) and the wild-type *NAM-B1*(1-215) sequence were cloned into the pDEST22 and pDEST32 vectors. These alleles consisted of the mutations present in the TILLING lines and the relevant alanine mutations, as described above (Supplementary Table 5, Additional File 2). Only the NAC domains were used to avoid auto-activation of the Y2H system caused by the transcriptional activation domain present in the C-terminal domain of the NAC transcription factors (23).

The *NAM-A1*(1-217) alleles in either vector were co-transformed with *NAM-B1*(1-215) in the alternative vector into chemically competent MaV203 cells. Simultaneously, individual vectors both with and without the *NAM-A1*(1-217) alleles were transformed into MaV203 as auto-activation controls. The transformations were recovered on SC-Leu-Trp media.

Initially, single colonies from the transformations were resuspended in 200 μL of dH_2_0. 2 μL of the colony suspension was plated on SC-Leu-Trp (control) and SC-Leu-Trp-His + 10μM 3AT (selection). These drop assays were scored for the presence of growth on the selection media as indicative of protein interactions. Six individual colonies were tested for each transformation. The experiment was repeated three times with consistent results.

Dilution assays were also carried out to highlight the relative strengths of the protein interactions. Here single colonies were grown overnight in a culture of liquid SC-Leu-Trp media to an OD of 0.1. From this, serial 1:10 dilutions were carried out in dH_2_0. Three individual colonies were tested for each combination of NAM-B1 in pDEST32 and the NAM-A1 allele in pDEST22. The experiment was repeated twice with consistent results.

#### Cell-death Scoring in *N. benthamiana*

Full-length *NAM-A1* alleles were cloned into the Gateway binary vector pGWB12, with an N-terminal FLAG tag (56). These were transformed into Agrobacterium (strain LBA4404) and co-infiltrated into 3-week-old *N. benthamiana* leaves alongside the silencing inhibitor P19 at a total OD of 0.4, and individual OD of 0.2 per construct. Leaves were harvested 3 days post-infiltration and snap-frozen in liquid N_2_. Leaf tissue was ground in liquid N_2_, protein was extracted as described above, and run on a 12% SDS-PAGE gel. The protein was transferred to a nitrocellulose membrane and probed with an anti-FLAG HRP-conjugated antibody (Abcam). Total protein input was visualised using a Ponceau stain.

The *NAM-A1* alleles were also patch-infiltrated into *N. benthamiana* leaves at the OD detailed above. The infiltrated patches were scored for cell death at 5 days post-infiltration, as detailed in Maqbool *et al.* 2015 (33). Briefly, images were taken of the abaxial and adaxial sides of the leaves using U.V. and white light, respectively. The infiltrated patches were then scored on a discrete scale from 0-6 based on the intensity of the fluorescence or chlorosis. Infiltrations were repeated on at least 20 independent leaves per allele in a randomised pattern around the leaf.

#### Data Analysis and Visualisation

All data analysis was carried out using R (v3.5.1). Graphs were produced using ggplot2 (57). Statistics were carried out using the base R package. Statistics tests used are indicated within the text where possible. Non-parametric statistical tests (two-sample Wilcoxon, Kruskal-Wallis Rank Sum) were used when assumptions of normality could not be met for analysis of glasshouse and field trial data. The Student’s t-test was used when normality assumptions were met. To account for the non-normal and discrete nature of the data in the *N. benthamiana* cell death experiment, statistical comparisons of cell death responses of the mutant alleles to wild-type *NAM-A1* were carried out by fitting a linear model using the Poisson distribution.

### Declarations

#### Ethics approval and consent to participate

Not applicable.

#### Consent for publication

Not applicable.

#### Availability of data and material

Original TILLING lines used in the experiment can be ordered from the SeedStor (https://www.seedstor.ac.uk/). All plasmids containing the wild-type *NAM-A1* and *NAM-B1* sequences used in this article have been deposited in Addgene.

#### Competing interests

The authors declare that they have no competing interests.

#### Funding

This work was funded by the UK Biotechnology and Biological Sciences Research Council (BBSRC) grants BB/P013511/1, BB/P016855/1 and Anniversary Future Leaders Fellowship to PB (BB/M014045/1). SAH was funded by the John Innes Foundation. LEO was supported by the BBSRC Norwich Research Park Biosciences Doctoral Training Partnership Research Experience Placement Scheme (grant number BB/M011216/1). NC was funded by Fulbright and CONICYT Becas-Chile 72111195.

#### Authors’ contributions

SAH, PB, and CU conceived the study. SAH and LEO cloned the NAM-A1 alleles and carried out the yeast 2-hybrid screen. SAH, PB, and NC carried out field trials. SAH carried out glasshouse trials, the *N. benthamiana* screen and the analysis of the conservation of the NAC domain. SAH and CU wrote the paper, and all authors contributed comments to the manuscript.

## Acknowledgements

The authors would like to thank J. Dubcovsky (UC Davis) for support during field trials in Davis, CA, and the glasshouse and field teams from the John Innes Centre and the University of California, Davis. They would also like to acknowledge M. Banfield and A. Smith (JIC) for useful comments during the development of this manuscript.

## Additional Files

**Additional File 1:** AdditionalFile1.clustal. “NAC domains from representative plant species”. This file contains the aligned NAC domains sequences from the seven plant species (*Triticum aestivum, Hordeum vulgare, Oryza sativa var. Japonica, Physcomitrella patens, Populus trichocarpa*, *Zea* mays, and *Arabidopsis thaliana*) that were included in the calculation of NAC domain conservation.

**Additional File 2:** AdditionalFile2.docx. “Supplementary Data”. This file contains supplementary figure and tables referred to in the text.

**Additional File 3:** AdditionalFile3.docx. “Gene and Protein Sequences of NAM genes”. This file contains the gene and protein sequences of the full and truncated version of NAM-A1 and NAM-B1 used for cloning and experiments.

