## Additional File 2 for "Conserved residues in the wheat (*Triticum aestivum*) NAM-A1 NAC domain are required for protein binding and when mutated lead to delayed peduncle and flag leaf senescence"

**Additional File 2: Supplementary Data**

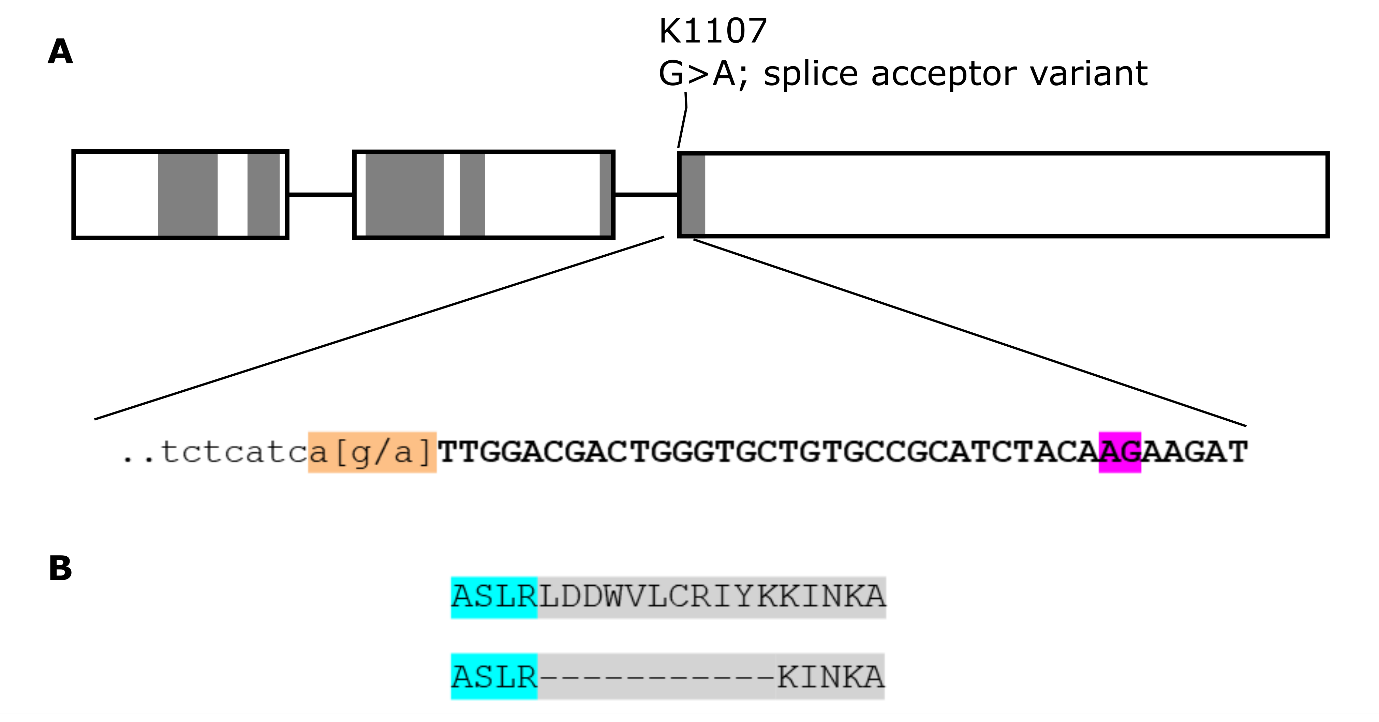

**Supplementary Figure 1: The splice acceptor variant in K1107 is predicted to lead to an 11-residue deletion.** The Kronos TILLING lines K1107 contains a G > A substitution at the splice acceptor site of the third exon (A). This mutation (orange) is predicted to lead to the use of an alternate splice site (purple). This predicted splice isoform contains an 11-residue deletion encompassing the majority of subdomain v (B).

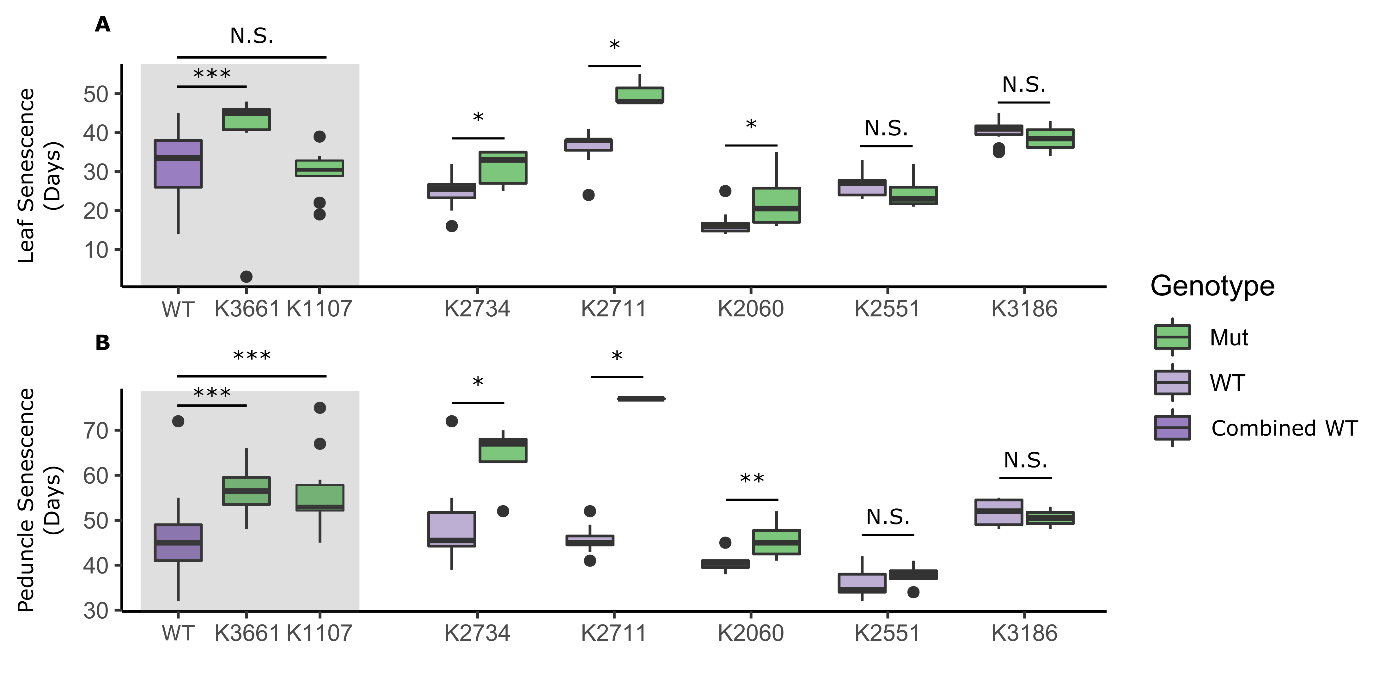

**Supplementary Figure 2: Lines K2711, K2734, and K3661 are consistently delayed in peduncle senescence.** Initial screening of the TILLING lines, excluding line K2615, found similar results for leaf senescence (A) and peduncle senescence (B). Notably, K2060 is also delayed in peduncle senescence in this experiment, and all lines containing missense mutations are slightly delayed in leaf senescence. The remaining mutant lines (K1107, K3661, and K2734) maintain the significant delay in peduncle senescence observed in the second experiment (Fig 2). The mutant plants for lines K3661 and K1107 are compared to the “Combined WT” of all WT plants from the remaining lines (see further details in methods). N is between 3 and 10. Statistical tests were carried out using the two-sample Wilcoxon test; *, p < 0.05; **, p < 0.01; ***, p < 0.001.

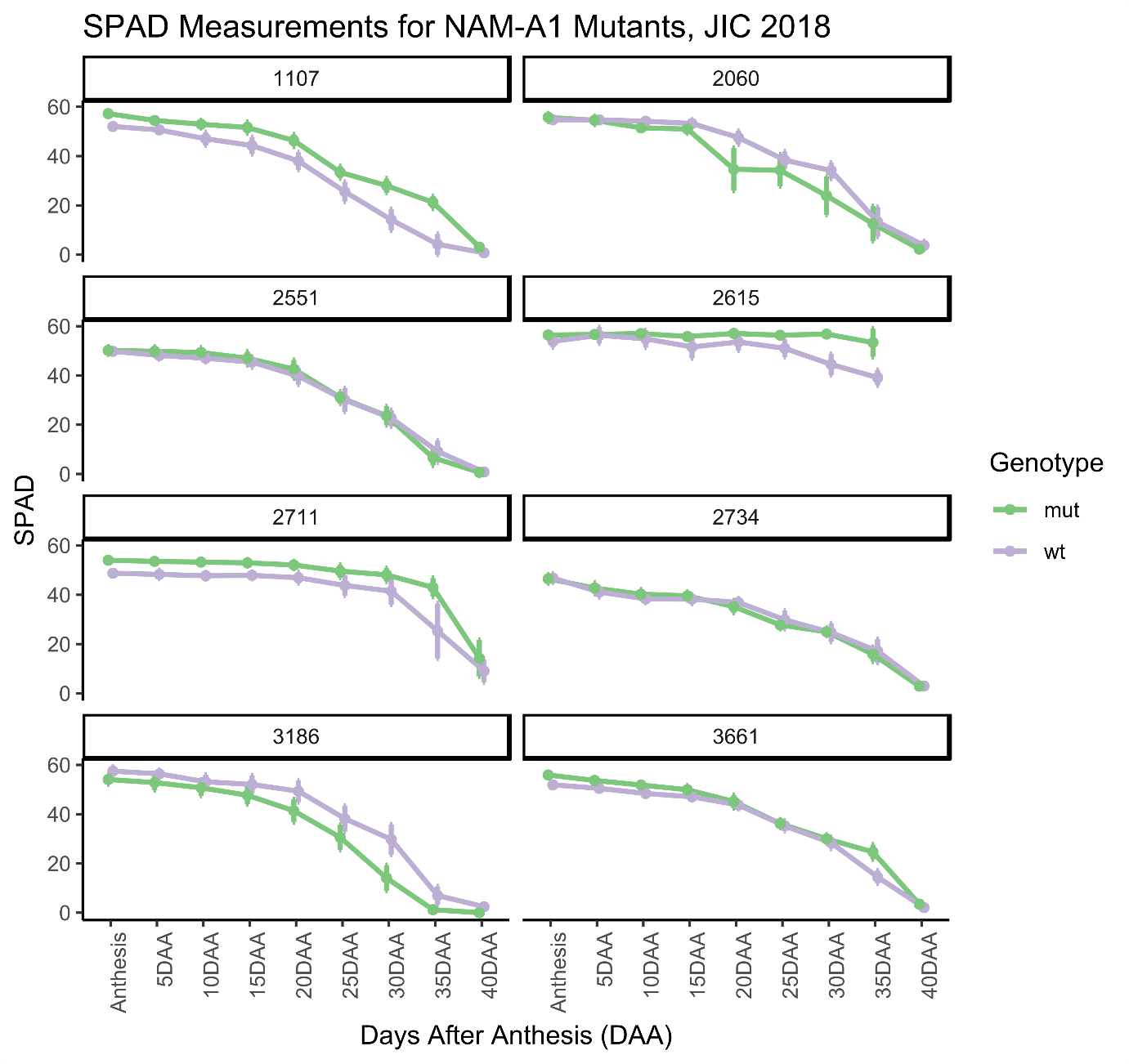

**Supplementary Figure 3: No significant differences in SPAD measurements are seen for the NAM-A1 TILLING mutants.** SPAD measurements were taken of the primary flag leaf from anthesis to 40 DAA in the glasshouse. Across all timepoints, no significant differences in SPAD units were observed for the mutant plants compared to wild-type for any of the TILLING lines. This correlates with no variation in leaf senescence onset being observed for the lines (Fig 2). N = 5 for all lines, genotypes, and timepoints. Statistical comparisons were carried out using the Kruskal-Wallis Rank Sum test.

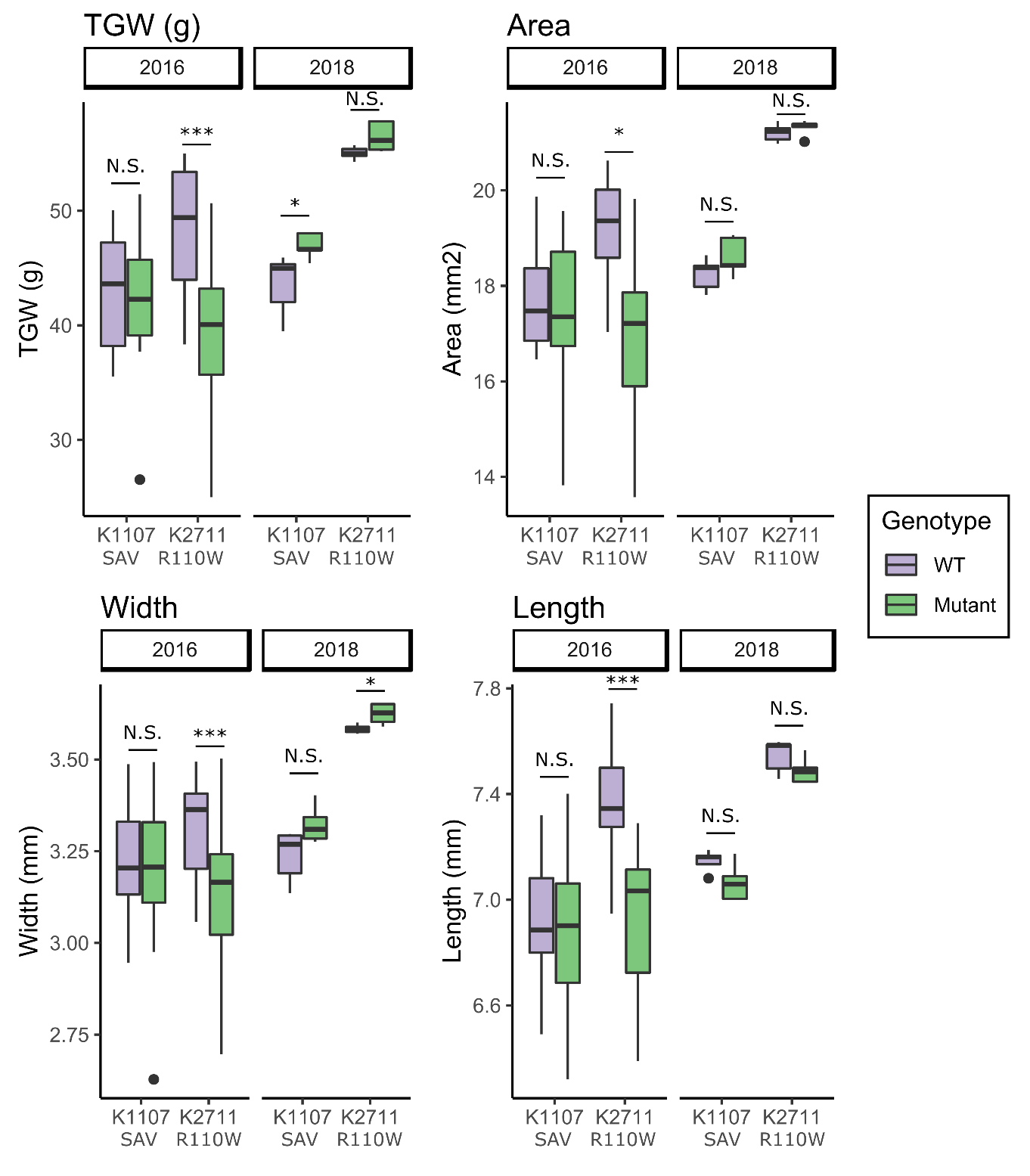

**Supplementary Figure 4: No consistent variation in grain size parameters is seen in the NAM-A1 mutant lines.** Across two years of field trials, neither the K2711 nor K1107 lines show any consistent variation in grain size or total grain weight (TGW). Variation in grain size and weight for K2711 in 2016 is most likely due to a background mutation co-segregating with the mutant allele that led to a substantial decrease in plant size (data not shown). This variation was lost in 2018, when lines had been selected to remove the co-segregating mutation. Statistical comparisons were carried out using the Kruskal-Wallis Rank Sum test; *, p < 0.05; **, p < 0.01; ***, p < 0.001.

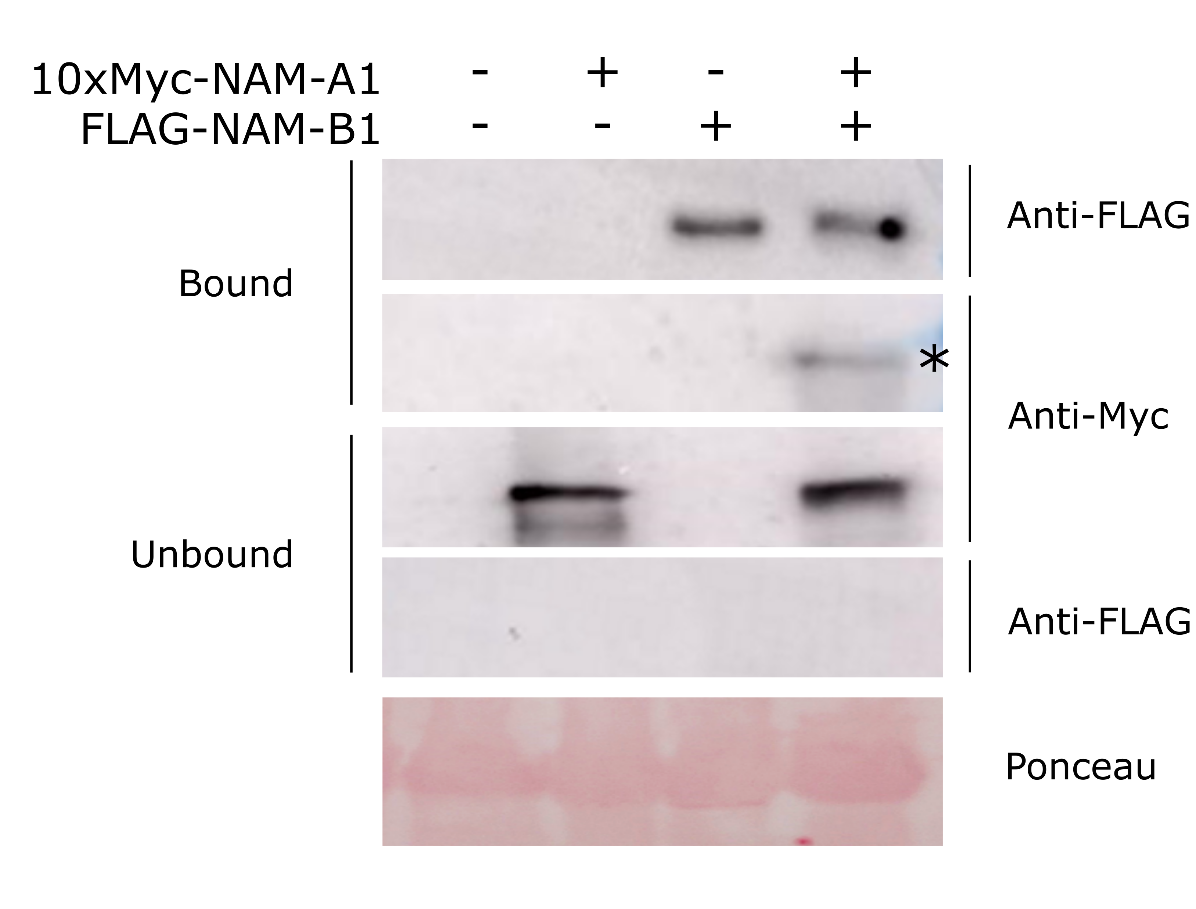

**Supplementary Figure 5: NAM-A1 interacts with NAM-B1 in a co-immunoprecipitation.** 10xMyc-tagged NAM-A1 is pulled down alongside FLAG-tagged NAM-B1 using anti-FLAG magnetic beads. FLAG-tagged bands are approximately 46 kDa; 10xMyc-tagged bands are approximately 51 kDa. Ponceau stain is of total protein input (Rubisco).

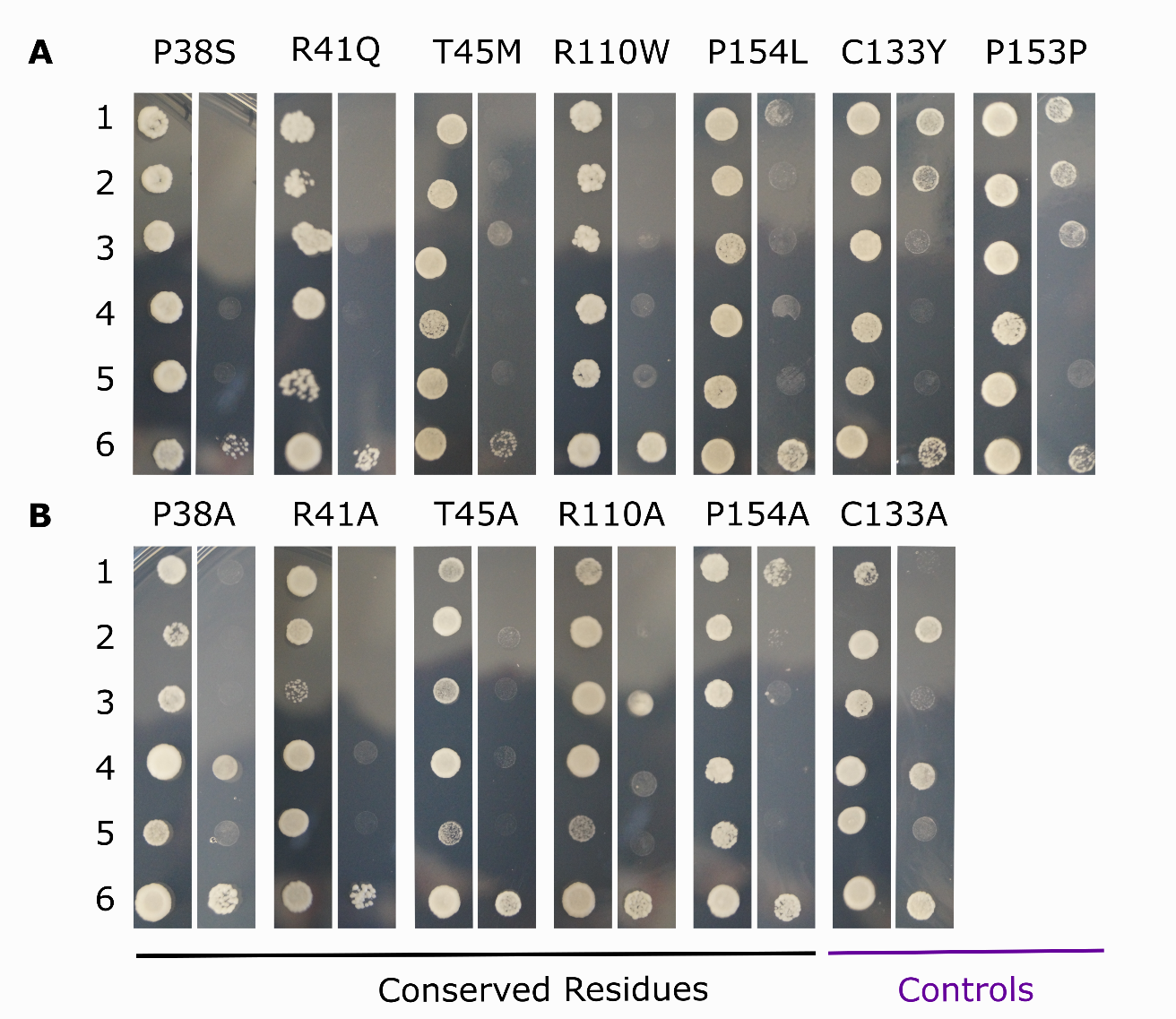

**Supplementary Figure 6: Mutations in conserved residues of NAM-A1 predominantly lead to loss of protein interaction.** TILLING (A) and alanine (B) mutant alleles of NAM-A1 were tested on control media (left; SC-LT) and selective media (right; SC-LTH + 10mM 3AT). NAM-A1 alleles were co-transformed with the wild-type NAM-B1 allele (1, NAM-A1 in pDEST22; 2, NAM-A1 in pDEST32). The ability of the mutant allele to interact with itself was also tested (3). Relevant controls were carried out for all alleles including NAM-A1:pDEST32 alone (4), NAM-A1:pDEST22 alone (5), and wild-type NAM-A1:pDEST22 with wild-type NAM-B1:pDEST32 (6).

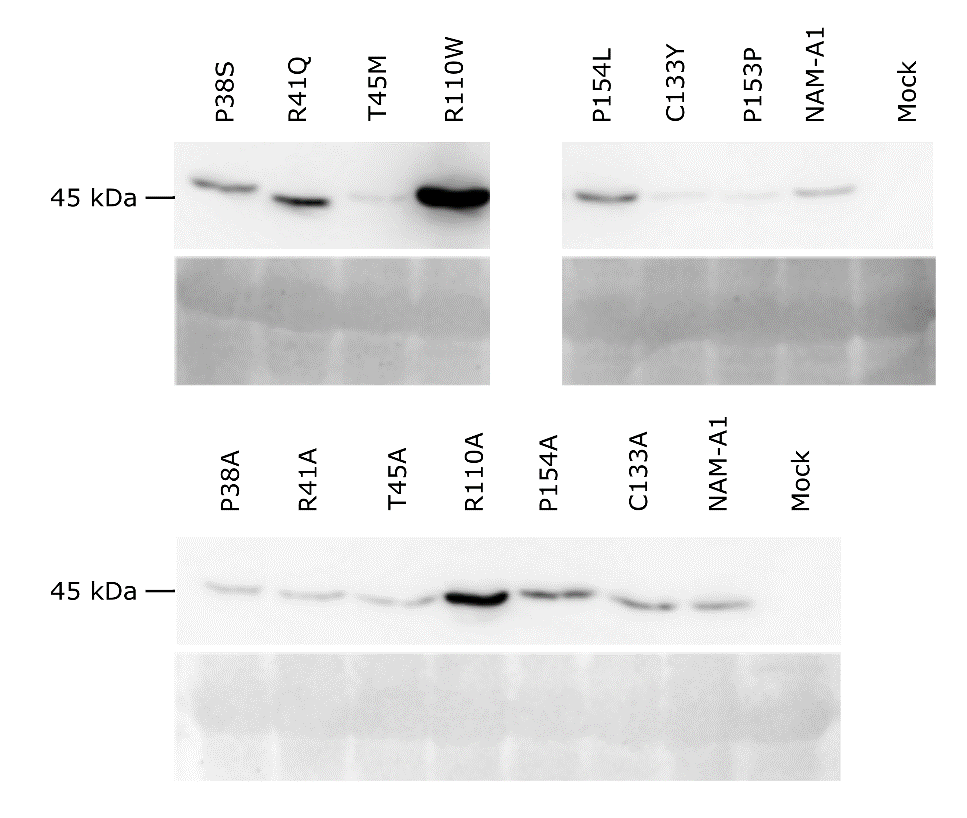

**Supplementary Figure 7: Expression levels of NAM-A1 mutant alleles does not correlate with observed HR response.** All constructs showed expression of the FLAG-tagged protein in *N. benthamiana* at 3 dpi. Expression levels were variable but did not correlate with observed cell death responses.

**Supplementary Tables**

| **Line** | **SNP** | **Codon mutation** | **Het/Hom** | **Consequence** | **cDNA position** | **CDS position** | **Amino acid mutation** | **SIFT** | **JSD Score** | **RefSeqv1 chr6 (bp)** | **Ensembl Link** |
| --- | --- | --- | --- | --- | --- | --- | --- | --- | --- | --- | --- |
| Kronos3661 | C/T | Ccg/Tcg | Hom | Missense | 339 | 112 | P38S | 0 | 0.26 | 77100016 | https://bit.ly/2GLlY71 |
| Kronos2615 | G/A | cGg/cAg | Het | Missense | 349 | 122 | R41Q | 0 | 0.4 | 77100006 | https://bit.ly/2X6gdX3 |
| Kronos2734 | C/T | aCg/aTg | Het | Missense | 361 | 134 | T45M | 0 | 0.46 | 77099994 | https://bit.ly/2S2xUTL |
| Kronos2711 | C/T | Cgg/Tgg | Het | Missense | 555 | 328 | R110W | 0 | 0.27 | 77099581 | https://bit.ly/2X1ZuUW |
| Kronos2551 | G/A | tGc/tAc | Het | Missense | 625 | 398 | C133Y | 0.78 | N/A | 77099511 | https://bit.ly/2Ig556S |
| Kronos2060 | C/T | cCc/cTc | Het | Missense | 688 | 461 | P154L | 0 | 0.17 | 77099448 | https://bit.ly/2SXh2lS |
| Kronos3186 | G/A | ccG/ccA | Het | Synonymous | 686 | 459 | P153P | N/A | 0.16 | 77099450 | https://bit.ly/2N6RCwK |
| Kronos1107 | G/A | N/A | Het | Splice Acceptor Variant | N/A | N/A | N/A | N/A | N/A | 77099212 | https://bit.ly/2GtGqtO |

**Supplementary Table 1: NAM-A1 mutation location and characterisation.** Details on the individual TILLING SNPs are provided, alongside the heterozygosity of the SNP in the M2 sequencing bulk, predicted consequence, and predicted impact based on conservation and SIFT scores. Links to the variant on EnsemblPlants are also provided.

**Supplementary Table 2: NAM genes are expressed in the peduncle and stem during reproduction.** Expression data derived from the Wheat Expression Browser (Borrill *et al.* 2016) was obtained for the five functional NAM copies (A1, A2, B2, D1, and D2) from the developmental time course of Azhurnaya (Ramirez-Gonzalez *et al.* 2018). Expression data is given in TPM, ± SEM.

|  |  | *NAM-A1* | *NAM-D1* | *NAM-A2* | *NAM-B2* | *NAM-D2* |
| --- | --- | --- | --- | --- | --- | --- |
| **Timepoint** | **Tissue** | TraesCS6A02G108300 | TraesCS6D02G096300 | TraesCS2A02G201800 | TraesCS2B02G228900 | TraesCS2D02G214100 |
| Ear emergence | Internode #2 | 1.47 ± 0.66 | 1.5 ± 0.92 | 1.26 ± 0.48 | 1.43 ± 0.94 | 1.84 ± 0.9 |
|  | Peduncle | 0.22 ± 0.06 | 0.49 ± 0.32 | 0.15 ± 0.17 | 0 | 0.01 ± 0.02 |
|  | Flag Leaf | 0.46 ± 0.05 | 0.13 ± 0.07 | 0.11 ± 0.11 | 1.19 ± 0.58 | 1.38 ± 0.05 |
| 30% spike | Internode #2 | 0.61 ± 0.37 | 0.55 ± 0.41 | 0.28 ± 0.16 | 0.25 ± 0.19 | 0.87 ± 0.85 |
|  | Peduncle | 0.24 ± 0.06 | 0 | 0.03 ± 0.03 | 0.01 ± 0.01 | 0 |
|  | Flag Leaf | 0.52 ± 0.07 | 0.06 ± 0.01 | 0.1 ± 0.12 | 0.37 ± 0.28 | 0.59 ± 0.28 |
| Milk grain | Internode #2 | 2.57 ± 0.08 | 3.05 ± 0.15 | 5.18 ± 0.96 | 5.42 ± 0.42 | 6.16 ± 0.74 |
|  | Peduncle | 3.56 ± 0.52 | 10.01 ± 1.5 | 11.51 ± 1.66 | 9.79 ± 1.26 | 5.22 ± 1.52 |
|  | Flag Leaf | 1.25 ± 0.46 | 0.3 ± 0.21 | 0.29 ± 0.37 | 4.17 ± 1.99 | 4.12 ± 1.26 |
| Dough | Flag Leaf | 2.92 ± 0.45 | 1.71 ± 0.67 | 1.55 ± 0.65 | 18.79 ± 7.47 | 13.61 ± 3.98 |
| Ripening | Flag Leaf | 9.59 ± 0.93 | 10.03 ± 1.65 | 8.04 ± 0.44 | 48.69 ± 10.66 | 30.25 ± 6.88 |

**Supplementary Table 3: KASP Primers for NAM-A1 TILLING mutants**

| **Line** | **WT (HEX/VIC)** | **Mut (FAM)** | **Common** |
| --- | --- | --- | --- |
| K3661 | GTGGAACCGGAAGCCCGG | GTGGAACCGGAAGCCCGA | GGCTCGGCGCAAAAAGCA |
| K2734 | CGACCAGCTCCTCGTCCG | CGACCAGCTCCTCGTCCA | AAGCAGCGCGGCATCAGCATG |
| K3741 | CTACTGGAAGGCCACCGG | CTACTGGAAGGCCACCGA | CGCCTTCTTGACGCCGAG |
| K2060 | TTTCTACCGCGGGAAGCCGCC | TTTCTACCGCGGGAAGCCGCT | TGGTGGTGGTGGAGCCAGACA |
| K3186 | TCTTCTACCGCGGGAAGCCG | TCTTCTACCGCGGGAAGCCA | TGGTGGTGGTGGAGCCAGACA |
| K2615 | GGAGCTCCCACCGGGCTTCCG | GGAGCTCCCACCGGGCTTCCA | AAGCAGCGCGGCATCAGCATG |
| K2551 | TGGCCTCGGGGACGGGGTG | TGGCCTCGGGGACGGGGTA | AAGCAGCGCGGCATCAGCATG |
| K2711 | AGCCCGACGTCGCCGCCCG | AGCCCGACGTCGCCGCCCA | TCCATCCATCAGAGAAGGCG |
| K1107 | AGCACCCAGTCGTCCAAC | AGCACCCAGTCGTCCAAT | TCGCACGGTATAGCTAGCAG |

**Supplementary Table 4: Primers used to amplify the full length and NAC domain fragments (“s”) of wild-type *NAM-A1* and *NAM-B1***

| **Construct** | **Forward** | **Reverse** | **Size of fragment (bp)** |
| --- | --- | --- | --- |
| *NAM-A1* | ATGAGGTCCATGGGCAG | TCAGGGATTCCAGTTCACG | 1227 |
| *NAM-B1* | ATGGGCAGCTCCGACTCA | CGTGAACTGGAATCCCTGA | 1218 |
| *NAM-A1s* | ATGAGGTCCATGGGCAG | TCACTGCTGATCTCCGG | 651 |
| *NAM-B1s* | ATGGGCAGCTCCGACTCA | CCGGCGATCAGCAGTGA | 645 |

**Supplementary Table 5: Primer sequences for site-directed mutagenesis of the *NAM-A1* alleles.**

|  | **ID** | **Line** | **SNP** | **Primer F** | **Primer R** | **Codon Change** | **Residue Change** |
| --- | --- | --- | --- | --- | --- | --- | --- |
| ***TILLING Mutants*** | SD1_P38S | Kronos3661 | C/T | CGGAGCTCCCATCGGGCTTCCGG | CCGGAAGCCCGATGGGAGCTCCG | Ccg/Tcg | P/S |
|  | SD1_R41Q | Kronos2615 | G /A | TCCCACCGGGCTTCCAGTTCCACCC | GGGTGGAACTGGAAGCCCGGTGGGA | cGg/cAg | R/Q |
|  | SD1_T45M | Kronos2734 | C /T | TTCCACCCGATGGACGAGGAGCTGGT | CGACCAGCTCCTCGTCCATCGGGTG | aCg/aTg | T/M |
|  | SD3_R110W | Kronos2711 | C/T | CGCGGCCGAACTGGGCGGCGACG | CGTCGCCGCCCAGTTCGGCCGCG | Cgg/Tgg | R/W |
|  | SD3_G121D | Kronos3741 | G /A | TACTGGAAGGCCACCGACACGGACAA | TTGTCCGTGTCGGTGGCCTTCCAGTA | gGc/gAc | G/D |
|  | SD4_P154L | Kronos2060 | C/T | AAGCCGCTCAAGGGCCTCAAAACCA | TGGTTTTGAGGCCCTTGAGCGGCTT | cCc/cTc | P/L |
|  | SD4_P153P | Kronos3186 | G/A | CGGGAAGCCGCCCAAGGGCC | TGAGGCCCTTGGGCGGCTTC | ccG/ccA | P |
|  | NC_C133Y | Kronos2551 | G/A | ACGGGGTACGGCCTGGTCCG | AGGCCGTACCCCGTCCCCGA | tGc/tAc | C/Y |
|  | **ID** | **Line** | **SNP** | **Primer F** | **Primer R** | **Codon Change** | **Residue Change** |
| ***Alanine Mutants*** | SD1_P38A | Kronos3661 | C/G | CGGAGCTCCCGTCGGGCTTCCGG | CCGGAAGCCCGCTGGGAGCTCCG | Ccg/Gcg | P/A |
|  | SD1_R41A | Kronos2615 | CG/GC | TCCCACCGGGCTTCGCGTTCCACCC | GGGTGGAACGCGAAGCCCGGTGGGA | CGg/GCg | R/A |
|  | SD1_T45A | Kronos2734 | A/G | TTCCACCCGGCGGACGAGGAGCTGGT | CGACCAGCTCCTCGTCCGCCGGGTG | Acg/Gcg | T/A |
|  | SD3_R110A | Kronos2711 | CG/GC | CGCGGCCGAACGCGGCGGCGACG | CGTCGCCGCCGCGTTCGGCCGCG | CGg/GCg | R/A |
|  | SD3_G121A | Kronos3741 | G/C | TACTGGAAGGCCACCGCCACGGACAA | TTGTCCGTGGCGGTGGCCTTCCAGTA | gGc/gCc | G/A |
|  | SD4_P154A | Kronos2060 | C/G | AAGCCGGCCAAGGGCCTCAAAACCA | TGGTTTTGAGGCCCTTGGCCGGCTT | Ccc/Gcc | P/A |
|  | NC_C133A | Kronos2551 | TG/GC | ACGGGGGCCGGCCTGGTCCG | AGGCCGGCCCCCGTCCCCGA | TGc/GCc | C/A |
