## Additional File 3 for "Conserved residues in the wheat (*Triticum aestivum*) NAM-A1 NAC domain are required for protein binding and when mutated lead to delayed peduncle and flag leaf senescence"

**Additional File 3**: CDS and protein sequences of *NAM-A1* and *NAM-B1* used as wild-type controls for yeast 2-hybrid and *N. benthamiana* cell death assays. Subdomains i – v are highlighted in green on the protein sequences.

> TraesCS6A01G108300.2; NAM-A1

CDS:

ATGAGGTCCATGGGCAGCTCCGACTCATCCTCCGGCTCGGCGCAAAAAGCAGCGCGGCATCAGCATGAGCCGCCGCCTCCGCGGCAGCGGGGCTCGGCGCCGGAGCTCCCACCGGGCTTCCGGTTCCACCCGACGGACGAGGAGCTGGTCGTGCACTACCTCAAGAAGAAGGCCGCCAAGGTGCCGCTCCCCGTCACCATCATCGCCGAGGTGGATCTCTACAAGTTCGACCCATGGGAGCTCCCCGAGAAGGCGACCTTCGGGGAGCAGGAGTGGTACTTCTTCAGCCCGCGCGACCGCAAGTACCCCAACGGCGCGCGGCCGAACCGGGCGGCGACGTCGGGCTACTGGAAGGCCACCGGCACGGACAAACCTATCCTGGCCTCGGGGACGGGGTGCGGCCTGGTCCGGGAGAAGCTCGGCGTCAAGAAGGCGCTCGTCTTCTACCGCGGGAAGCCGCCCAAGGGCCTCAAAACCAACTGGATCATGCACGAGTACCGCCTCACCGACGCGTCTGGCTCCACCACCACCAGCCGGCCGCCGCCGCCTGTGACCGGCGGGAGCCGGGCTGCAGCCTCTCTGAGGTTGGACGACTGGGTGCTGTGCCGCATCTACAAGAAGATCAACAAGGCCGCGGCCGGAGATCAGCAGAGGAGCACGGAGTGCGAGGACTCCGTGGAGGACGCGGTCACCGCGTACCCGCTCTATGCCACGGCGGGCATGGCCGGTGCAGGTGCGCATGGCAGCAACTACGCTTCACCTTCACTGCTCCATCATCAGGACAGCCATTTCCTGGAGGGCCTGTTCACAGCAGACGACGCCGGCCTCTCGGCGGGCGCCACCTCGCTGAGCCACCTGGCCGCGGCGGCGAGGGCGAGCCCGGCTCCGACCAAACAGTTTCTCGCCCCGTCGTCTTCAACCCCGTTCAACTGGCTCGATGCGTCACCCGCCGGCATCCTGCCACAGGCAAGGAATTTCCCTGGGTTTAACAGGAGCAGAAACGTCGGCAATATGTCGCTGTCATCGACGGCCGACATGGCTGGCGCGGCCGGCAATGCGGTGAACGCCATGTCCGCATTTATGAATCCTCTCCCCGTGCAAGACGGGACGTACCATCAACACCATGTCATCCTCGGCGCCCCACTGGCGCCAGAGGCTACCACAGGCGGCGCCACCTCTGGTTTCCAGCATCCCGTCCAAGTATCCGGCGTGAACTGGAATCCCTGA

Protein:

MRSMGSSDSSSGSAQKAARHQHEPPPPRQRGSAPELPPGFRFHPTDEELVVHYLKKKAAKVPLPVTIIAEVDLYKFDPWELPEKATFGEQEWYFFSPRDRKYPNGARPNRAATSGYWKATGTDKPILASGTGCGLVREKLGVKKALVFYRGKPPKGLKTNWIMHEYRLTDASGSTTTSRPPPPVTGGSRAAASLRLDDWVLCRIYKKINKAAAGDQQRSTECEDSVEDAVTAYPLYATAGMAGAGAHGSNYASPSLLHHQDSHFLEGLFTADDAGLSAGATSLSHLAAAARASPAPTKQFLAPSSSTPFNWLDASPAGILPQARNFPGFNRSRNVGNMSLSSTADMAGAAGNAVNAMSAFMNPLPVQDGTYHQHHVILGAPLAPEATTGGATSGFQHPVQVSGVNWNP*

> TraesCS6A01G108300.2; NAM-A1 (NAC domain, 1-217)

CDS:

ATGAGGTCCATGGGCAGCTCCGACTCATCCTCCGGCTCGGCGCAAAAAGCAGCGCGGCATCAGCATGAGCCGCCGCCTCCGCGGCAGCGGGGCTCGGCGCCGGAGCTCCCACCGGGCTTCCGGTTCCACCCGACGGACGAGGAGCTGGTCGTGCACTACCTCAAGAAGAAGGCCGCCAAGGTGCCGCTCCCCGTCACCATCATCGCCGAGGTGGATCTCTACAAGTTCGACCCATGGGAGCTCCCCGAGAAGGCGACCTTCGGGGAGCAGGAGTGGTACTTCTTCAGCCCGCGCGACCGCAAGTACCCCAACGGCGCGCGGCCGAACCGGGCGGCGACGTCGGGCTACTGGAAGGCCACCGGCACGGACAAACCTATCCTGGCCTCGGGGACGGGGTGCGGCCTGGTCCGGGAGAAGCTCGGCGTCAAGAAGGCGCTCGTCTTCTACCGCGGGAAGCCGCCCAAGGGCCTCAAAACCAACTGGATCATGCACGAGTACCGCCTCACCGACGCGTCTGGCTCCACCACCACCAGCCGGCCGCCGCCGCCTGTGACCGGCGGGAGCCGGGCTGCAGCCTCTCTGAGGTTGGACGACTGGGTGCTGTGCCGCATCTACAAGAAGATCAACAAGGCCGCGGCCGGAGATCAGCAG

Protein:

MRSMGSSDSSSGSAQKAARHQHEPPPPRQRGSAPELPPGFRFHPTDEELVVHYLKKKAAKVPLPVTIIAEVDLYKFDPWELPEKATFGEQEWYFFSPRDRKYPNGARPNRAATSGYWKATGTDKPILASGTGCGLVREKLGVKKALVFYRGKPPKGLKTNWIMHEYRLTDASGSTTTSRPPPPVTGGSRAAASLRLDDWVLCRIYKKINKAAAGDQQ*

> DQ869673.1; NAM-B1

CDS:

ATGGGCAGCTCCGACTCATCTTCCGGCTCGGCGCAAAAAGCAACGCGGTATCACCATCAGCATCAGCCGCCGCCTCCGCAGCGGGGCTCGGCGCCGGAGCTCCCGCCGGGCTTCCGGTTCCACCCGACGGACGAGGAGCTGGTGGTGCACTACCTCAAGAAGAAGGCCGACAAGGCGCCGCTCCCCGTCAACATCATCGCCGAGGTGGATCTCTACAAGTTCGACCCATGGGAGCTCCCCGAGAAGGCGACCATCGGGGAGCAGGAGTGGTACTTCTTCAGCCCGCGCGACCGCAAGTACCCCAACGGCGCGCGGCCGAACCGGGCGGCGACGTCGGGGTACTGGAAGGCCACCGGCACGGACAAGCCTATCCTGGCCTCGGGGACGGGGTGCGGCCTGGTCCGGGAGAAGCTCGGCGTCAAGAAGGCGCTCGTGTTCTACCGCGGGAAGCCGCCCAAGGGCCTCAAAACCAACTGGATCATGCACGAGTACCGCCTCACCGACGCATCTGGCTCCACCACCGCCACCAACCGACCGCCGCCGGTGACCGGCGGGAGCAGGGCTGCTGCCTCTCTCAGGTTGGACGACTGGGTGCTGTGCCGCATCTACAAGAAGATCAACAAGGCCGCGGCCGGCGATCAGCAGAGGAACACGGAGTGCGAGGACTCCGTGGAGGACGCGGTCACCGCGTACCCGCTCTATGCCACGGCGGGCATGACCGGTGCAGGTGCGCATGGCAGCAACTACGCTTCACCTTCACTGCTCCATCATCAGGACAGCCATTTCCTGGACGGCCTGTTCACAGCAGACGACGCCGGCCTCTCGGCGGGCGCCACCTCGCTGAGCCACCTAGCAGCGGCGGCGAGGGCAAGCCCGGCTCCGACCAAACAGTTTCTCGCCCCGTCGTCTTCGACCCCGTTCAACTGGCTCGATGCGTCACCAGTCGGCATCCTCCCACAGGCAAGGAATTTTCCTGGGTTTAACAGGAGCAGAAACGTCGGCAATATGTCGCTGTCATCGACGGCCGACATGGCTGGCGCAGTGGACAACGGTGGAGGCAATGCGGTGAACGCCATGTCTACCTATCTTCCCGTGCAAGACGGGACGTACCATCAGCAGCATGTCATCCTCGGCGCTCCGCTGGTGCCAGAAGCCGCCGCCGCCACCTCTGGATTCCAGCATCCCGTTCAAATATCCGGCGTGAACTGGAATCCCTGA

Protein:

MGSSDSSSGSAQKATRYHHQHQPPPPQRGSAPELPPGFRFHPTDEELVVHYLKKKADKAPLPVNIIAEVDLYKFDPWELPEKATIGEQEWYFFSPRDRKYPNGARPNRAATSGYWKATGTDKPILASGTGCGLVREKLGVKKALVFYRGKPPKGLKTNWIMHEYRLTDASGSTTATNRPPPVTGGSRAAASLRLDDWVLCRIYKKINKAAAGDQQRNTECEDSVEDAVTAYPLYATAGMTGAGAHGSNYASPSLLHHQDSHFLDGLFTADDAGLSAGATSLSHLAAAARASPAPTKQFLAPSSSTPFNWLDASPVGILPQARNFPGFNRSRNVGNMSLSSTADMAGAVDNGGGNAVNAMSTYLPVQDGTYHQQHVILGAPLVPEAAAATSGFQHPVQISGVNWNP*

> DQ869673.1; NAM-B1 (NAC domain, 1-215)

CDS:

ATGGGCAGCTCCGACTCATCTTCCGGCTCGGCGCAAAAAGCAACGCGGTATCACCATCAGCATCAGCCGCCGCCTCCGCAGCGGGGCTCGGCGCCGGAGCTCCCGCCGGGCTTCCGGTTCCACCCGACGGACGAGGAGCTGGTGGTGCACTACCTCAAGAAGAAGGCCGACAAGGCGCCGCTCCCCGTCAACATCATCGCCGAGGTGGATCTCTACAAGTTCGACCCATGGGAGCTCCCCGAGAAGGCGACCATCGGGGAGCAGGAGTGGTACTTCTTCAGCCCGCGCGACCGCAAGTACCCCAACGGCGCGCGGCCGAACCGGGCGGCGACGTCGGGGTACTGGAAGGCCACCGGCACGGACAAGCCTATCCTGGCCTCGGGGACGGGGTGCGGCCTGGTCCGGGAGAAGCTCGGCGTCAAGAAGGCGCTCGTGTTCTACCGCGGGAAGCCGCCCAAGGGCCTCAAAACCAACTGGATCATGCACGAGTACCGCCTCACCGACGCATCTGGCTCCACCACCGCCACCAACCGACCGCCGCCGGTGACCGGCGGGAGCAGGGCTGCTGCCTCTCTCAGGTTGGACGACTGGGTGCTGTGCCGCATCTACAAGAAGATCAACAAGGCCGCGGCCGGCGATCAGCAGTGA

Protein:

MGSSDSSSGSAQKATRYHHQHQPPPPQRGSAPELPPGFRFHPTDEELVVHYLKKKADKAPLPVNIIAEVDLYKFDPWELPEKATIGEQEWYFFSPRDRKYPNGARPNRAATSGYWKATGTDKPILASGTGCGLVREKLGVKKALVFYRGKPPKGLKTNWIMHEYRLTDASGSTTATNRPPPVTGGSRAAASLRLDDWVLCRIYKKINKAAAGDQQ*
